# Structures of the cyanobacterial nitrogen regulators NtcA and PipX complexed to DNA fully clarify DNA binding by NtcA and recruitment of RNA polymerase by PipX

**DOI:** 10.1101/2024.08.29.607757

**Authors:** Alicia Forcada-Nadal, Sirine Bibak, Paloma Salinas, Asunción Contreras, Vicente Rubio, José L. Llácer

**Affiliations:** Instituto de Biomedicina de Valencia of the CSIC (IBV-CSIC), Valencia Spain; Group 739 at the IBV-CSIC of the Centro de Investigación Biomédica en Red en Enfermedades Raras of the Instituto de Salud Carlos III (CIBERER-ISCIII), Madrid, Spain; Departamento de Fisiología, Genética y Microbiología, Universidad de Alicante, Alicante, Spain

**Keywords:** Transcriptional regulation, PipX, NtcA, CRP, FNR, RNA polymerase, nitrogen regulation, cyanobacteria, *Synechococcus elongatus*

## Abstract

The CRP-FNR superfamily of transcriptional regulators includes the cyanobacterial master regulator NtcA, which orchestrates large responses of cyanobacteria to nitrogen scarcity. NtcA uses as allosteric activator 2-oxoglutarate (2OG), a signal of nitrogen poorness and carbon richness, and binds a coactivating protein (PipX) that shuttles between the signaling protein PII and NtcA depending on nitrogen richness, thus connecting PII signaling and gene expression regulation. Here, combining structural (X-ray crystallography of six types of crystals including NtcA complexes with DNA, 2OG and PipX), modelling and functional (EMSA and bacterial two-hybrid) studies, we clarify the reasons for the exquisite specificity for the binding of NtcA to its target DNA, its mechanisms of activation by 2OG, and its coactivation by PipX. Our crystal structures of PipX-NtcA-DNA complexes prove that PipX does not interact with DNA, although it increases NtcA-DNA contacts, and that it stabilizes the active, DNA-binding-competent conformation of NtcA. Superimposition of this complex on a very recently reported cryoEM structure of NtcA in a Transcription Activity Complex with RNA polymerase (RNAP), shows that PipX binding helps recruit RNAP by PipX interaction with RNAP, particularly with its gamma and sigma (region 4) subunits, a structural prediction supported here by bacterial two-hybrid experiments.

## Introduction

Regulation of gene expression is a crucial process for adaptation of bacteria to changing environments, colonization of diverse niches and pathogenicity. Key players in those processes are transcriptional regulators. Of paramount importance among transcriptional regulators is the superfamily of these proteins known as CRP-FNR (<u>c</u>AMP receptor <u>p</u>rotein-fumarate/<u>n</u>itrate reductase regulator), which encompasses at least 21 subclasses and is represented in most bacterial phyla (Körner *et al*. 2003; Krol *et al*. 2023). The members of this superfamily are homodimers composed of an N-terminal dimerization and effector-binding domain (EBD) and of a C-terminal DNA binding domain (DBD) with a helix-turn-helix (HTH) motif. Members of this superfamily regulate a broad range of processes, including responses to hunger, oxygen, carbon monoxide, oxidative stress, nitrogen fixation, availability to metabolically active nitrogen forms and photosynthesis (Körner *et al*. 2003; Roberts 2024; Vegas-Palas *et al*. 1992).

CRP is considered the paradigm for this superfamily and has reached textbook-example status (Lawson *et al* 2004). The determination, more than 30 years ago, of the crystal structure of CRP bound to its target DNA revealed determinants involved in the specificity of DNA sequence recognition (Schultz *et al*. 1991). DNA binding involved the insertion of the αF helices of the HTH motif from each subunit of the CRP dimer into adjacent major grooves of the double-stranded DNA. CRP binding induced a significant bend on the DNA, which was found to facilitate transcription activation (Shultz *et al*. 1991, Parkinson *et al*. 1996, Chen *et al*. 2001, Chen *et al*. 2001, Napoli *et al*. 2006, Benoff *et al*. 2002).

DNA binding and transcriptional regulation mechanisms appear to be shared by the members of the CRP-FNR superfamily. However, structural snapshots for protein-DNA complexes have been obtained in just a few cases (Levy *et al*. 2008; Hall *et al*. 2016; Bonnet *et al*. 2013; Werel *et al*. 2023) and there is even less information concerning transcription activation complexes (TACs), which include the RNA polymerase (RNAP) holoenzyme and is critical for understanding transcriptional regulation. A crystal structure of the bacterial class II TAC from *Thermus thermophilus* revealed how the CRP/FNR activator TAP enhances transcription initiation through stabilizing interactions with RNAP (Feng *et al*. 2016). Cryo-electron microscopy studies of *E. coli* CRP-TAC complexes of class I (Liu *et al*. 2017) and class II (Shi *et al*. 2020) types have illustrated diverse interactions between CRP and RNAP, offering detailed views of how CRP binding to DNA facilitates transcription by RNAP. Very recently, cryo-electron microscopy structures of *Anabaena* sp. PCC7120 TAC with the NtcA master regulator of cyanobacteria (a CRP/FNR superfamily member) have provided additional information on relevant interactions and on cooperative transcription activation by NtcA and the LysR-like regulator NtcB (Han *et al*. 2024).

NtcA, a central subject of the present work, is a global nitrogen regulator conserved in cyanobacteria that controls a very large regulon mainly involved in nitrogen assimilation (Vegas-Palas *et al*. 1992; Herrero *et al*. 2001; Herrero *et al*. 2004, Su *et al*. 2005; Mitschke *et al*. 2011; Espinosa *et al*. 2014; Giner-Lamia *et al*. 2017). NtcA binds 2-oxoglutarate (2OG), a small molecule that signals the intracellular carbon/nitrogen balance (high levels when carbon is abundant and there is poorness of metabolizable nitrogen; Llácer *et al*. 2008) and induces the active conformation of NtcA, binding at a topologically equivalent site to that of cAMP in CRP (Llácer *et al*. 2010; Zhao *et al*. 2010). However, NtcA is unique within the CRP/FNR family because it is coactivated by PipX, a small protein conserved in cyanobacteria (Burillo *et al*. 2004, Labella *et al*. 2020) that shuttles between the carbon/nitrogen/energy sensor/signaling protein PII and NtcA, depending on whether nitrogen richness is high or low (Espinosa *et al*. 2006, Labella *et al* 2016; Labella *et al*. 2017; Forcada-Nadal *et al*. 2018), thus enabling control by PII of gene expression in cyanobacteria (Llácer *et al*. 2010).

We previously (Llácer *et al*. 2010) determined a crystal structure of NtcA bound to PipX and proposed that PipX would stabilize the NtcA conformation that is competent for binding to target DNA (NtcA box). This box is an imperfect palindrome with three symmetrical specificity-conferring base pairs located 5-7 nucleotides upstream and downstream from the palindrome centre (Luque *et al*. 1994). Unfortunately, we were unable to obtain an experimental structure of the NtcA-DNA complex, and, therefore, this proposal was speculative, as it was, too, our proposal that PipX, by interacting with RNAP, could help recruit this polymerase to the NtcA binding site in DNA (Llácer *et al*. 2010). The recent structures of the NtcA TAC at the nirA promoter (Han *et al*. 2024) offer significant insights but provide no information on NtcA binding specificity or on the role of PipX.

Here we define the structural determinants of NtcA binding to target DNA by providing a 3 Å crystallographic structure of the NtcA of *Synechococcus elongatus* strain PCC7942 (from now on *S. elongatus*) bound to both DNA and 2OG. We also present a crystal structure of the quaternary complex of NtcA with PipX, 2OG and DNA, showing that PipX, without contacting promoter DNA sequences, substantially enhances the binding of NtcA to its target site. We also confirm our previous proposal that PipX binding freezes NtcA in its DNA binding-competent conformation. Importantly, by modelling our PipX-NtcA-DNA complex onto the NtcA-TAC structure (PDB 8H40) from Han et al. (2024) (a structure that did not contain PipX), and by subsequently testing the interactions between PipX and relevant domains of the sigma and gamma subunits of RNAP in the bacterial two-hybrid system, we strongly support that, indeed, PipX enhances NtcA-specific transcription by contributing to the recruitment of the RNAP to the NtcA binding sites in DNA.

Thus, our present work, in addition to defining in great structural detail the determinants of promoter selectivity of NtcA, expands the repertoire of mechanisms of transcriptional regulation, calling attention to the PipX coactivator protein, a specific regulator of cyanobacteria with no known functional counterparts in other taxonomic groups.

## Materials and Methods

### Production and crystallization of NtcA-DNA, PipX-NtcA-DNA and of NtcA in inactive apo forms

NtcA and PipX from *Synechococcus elongatus* (strain PCC7942) with N-terminal His6 tags were produced in *E. coli* Rosetta (DE3) cells (from Novagen) from plasmids pTrc99A-*pipX* and pET15b-NtcA, respectively, and were purified essentially as reported (Espinosa *et al*. 2006, Llácer *et al*. 2010). To produce the V187E mutant form of NtcA, site-directed mutagenesis of the pET15b-NtcA plasmid was carried out using the Quickchange II system (from Stratagene, La Jolla, CA), following the manufacturer’s instructions, and utilizing the oligonucleotide pair 5’TCGGCTCAACGCGGGAGACAGTGACG3’ and 5’CGTCACTGTCTCCCGCGTTGAGCCGATCG3’. After corroborating by DNA sequencing the introduction of the mutation, the NtcA mutant was expressed and purified as the corresponding wild-type protein.

The DNA used in the crystallization of the NtcA-2OG-DNA complex was prepared by 10-min heating at 65°C of an equimolar mixture of the synthetic oligodesoxyribonucleotides 5’-AGCTGA**TAC**ATAAAAAT-3’ and 5’-CATTTTTAT**GTA**TC-3’ (from Sigma) in 5 mM Tris-HCl, pH 7.5, followed by slow cooling to 4°C for annealing. These oligonucleotides include one half of the NtcA binding site of the *S. elongatus glnA* promoter (consensus bases are in bold type), to produce upon annealing the following chained duplex (underlining and italics are used to differentiate individual oligonucleotides; bold-type, consensus bases in the NtcA box): 5’<u>CATTTTTAT**GTA**TC</u>*AGCTGA**TAC**ATAAAAAT 3’* 3’<u>TAAAAATA**CAT**AGTCGA</u>*CT**ATG**TATTTTTAC 5’*

Then, 0.3 mg/ml of this annealed DNA mixture and 0.85 mg/ml NtcA in buffer A (50 mM Na citrate, pH 6.5, 0.5 M NaCl, 5 mM MgCl2, 50 mM L-arginine-HCl, 50 mM Na L-glutamate and 10 mM 2OG), was incubated 5 min at 22°C, and concentrated to 7.3 mg protein/ml by ultrafiltration in an Amicon Ultracel YM-50 device. Crystallization sitting drops were prepared by mixing 0.4 μl of the protein-DNA solution and 0.4 μl of crystallization solution. The crystal used was obtained at 21°C in a crystallization solution consisting of 0.1 M Bis-Tris, pH 6.5 and 28% polyethylene glycol (PEG) monomethyl ether 2K.

The procedure to crystallize the PipX-2OG-NtcA-DNA complex was similar to that for the NtcA-DNA complex, although the annealed DNA contained the whole perfect NtcA box (consensus bases are in bold type) in a symmetric non-cleaved 30-bp DNA duplex with sticky ends (5’ protruding C and 3’ overhanging G in one and the other oligonucleotides) to create the following duplex: 5’CATTTTTAT**GTA**TCAGCTGA**TAC**ATAAAAAT TAAAAATA**CAT**AGTCGACT**ATG**TATTTTTAG5’

We first prepared the NtcA-DNA complex by incubating 10 min at 22°C 0.6 mg/ml of this annealed DNA and 1.4 mg/ml NtcA in buffer A. Then 0.6 mg/ml PipX was added, the mixture was incubated 10 more min and then it was concentrated to ∼6 mg protein/ml by centrifugal ultrafiltration. Crystallization sitting drops were prepared as above, obtaining after several weeks two types of crystals with the following crystallization solutions: for crystal type I, 0.2 M MgSO4 and 20% PEG 3350; and for crystal type II 0.1 M Tris-HCl, pH 8.5, 2 M ammonium phosphate and 0.1 M ZnCl2. The first crystals obtained of this complex were of type II. They diffracted X-rays at 6 Å-resolution, with 63% solvent content. The resolution was improved to 4.3 Å in one crystal by controlled dehydration for 30 min via vapor diffusion versus 50 μl of a solution of 4 M KCl. Initial phases were obtained with this crystal (see below). Then we obtained crystals of type I that diffracted X-rays up to 3.8 Å resolution.

Crystals of NtcA without 2OG or DNA were obtained in three different forms from a 6.1 mg/ml NtcA solution prepared as above except for the omission of 2OG from all solutions and from buffer A, using as crystallization solutions, for form A1, 0.1 M Na acetate pH 4.5, 40% PEG 200; for form A2, 0.1 M Na HEPES pH 6.5, 20.5% PEG 4 K; and for form B1, 0.05 M Tris-HCl pH 8, 2 M NaCl, 2% 2-methyl-2,4-pentanediol, 10 mM MgCl2 and 5% DMSO.

### Data collection and structure determination

Crystals were flash-frozen in liquid N2. Cryoprotection of crystal II of PipX-NtcA-DNA was as indicated in the previous section. The crystal of NtcA apo form A1 did not need added cryoprotection. All other crystals were cryoprotected by harvesting them in their corresponding crystallization solutions fortified as indicated: for NtcA-DNA, 38% PEG monomethyl ether 2K; for type I crystals of PipX-NtcA-DNA, 39% PEG monomethyl ether 3.35K; and for NtcA apo forms A2 and B1, 25 % PEG 400, and 27% 2-methyl-2,4-pentanediol, respectively.

Diffraction was at 100 K in the indicated beamlines (Table 1) of the ESRF or Diamond synchrotrons. Datasets obtained for NtcA A1 and B1, at respective resolutions of 2.70 and 3.33 Å, and for PipX-NtcA-DNA I and II at 3.8 and 4.3 Å resolutions, respectively, were processed and scaled with MOSFLM and SCALA [CCP4 suite (Collaborative Computational Project Number 4, 1994)], whereas those for NtcA-DNA (3.00 Å resolution) and NtcA apo A2 (2.85 Å) were processed with XDS (Kabsch *et al*. 2010), and prepared with COMBAT (Collaborative Computational Project Number 4, 1994) for scaling with SCALA. See Table 1 for the space groups and cell dimensions for the different crystals.

**Table 1.**
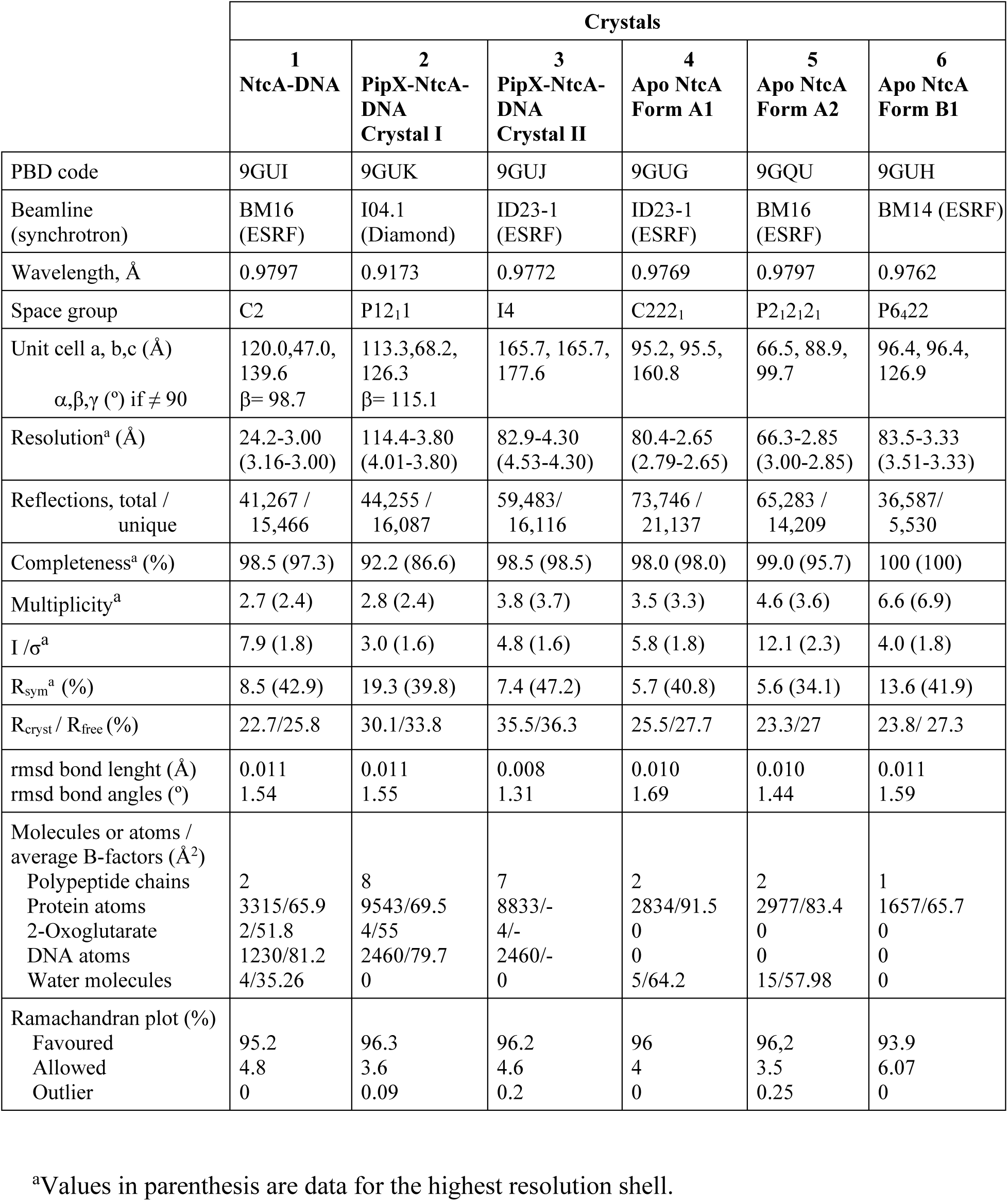
Data collection and refinement statistics.

Crystallographic phases for the NtcA-DNA crystal (containing also one bound molecule of 2OG per subunit) were determined by molecular replacement with MOLREP (Vagin *et al*. 2010) using as a model the *S. elongatus* NtcA dimer in its active conformation (PDB accession number 2XHK; Llácer *et. al.* 2010). The initial model was improved by rigid body and maximum likelihood restrained refinement using REFMAC (Murshudov *et al*. 2011). The phases obtained from this partially refined solution were improved by density modification using program PARROT (Cowtan, K. 2010). An initial model for DNA was manually built using COOT (Emsley *et al*. 2010). Then, iterative rounds of model building with COOT and refinement with REFMAC and PHENIX (Afonine *et al*. 2018) were carried out. Tight non-crystallographic symmetry (NCS) restraints were applied until the final rounds of refinement, when they were gradually released. TLS (Winn *et al*. 2001) was used in the last steps of refinement, defining the TLS groups after TLSMD analysis (Painter *et al*. 2006). All the diffraction data were used throughout the refinement processes, except the 5% randomly selected data for calculating *R*free. The stereochemical quality of the model was analysed with MolProbity (Williams *et al*. 2018). The final refined model, at 3.00 Å-resolution, contained one NtcA dimer and two chains of each oligodeoxyribonucleotide (Table 1). 2OG was modeled in the NtcA-DNA structure as seen in the active *Anabaena* structure (Zhao *et al*. 2010)

One subunit of the structure of active NtcA from *S. elongatus* was also used as a model for phasing the dataset for NtcA A1, by molecular replacement with MOLREP. Model building with COOT and refinement with REFMAC and TLS were as for the NtcA-DNA complex. One of the subunits of this final model (which contained two subunits in the asymmetric unit) was used for phasing the dataset for NtcA A2 by molecular replacement using PHASER (McCoy *et al*. 2007). A satisfactory solution was found, consisting in one dimer in the asymmetric unit. Model building and refinement were as above to a final resolution of 2.85 Å (Table 1). For NtcA B1, molecular replacement with MOLREP using a subunit of the inactive conformation found previously with NtcA from *S. elongatus* (PDB accession number 2XKP, Llácer *et al*. 2010) yielded a satisfactory solution, consisting in one subunit in the asymmetric unit, which was refined as with the other datasets, generating a dimer by application of the crystal symmetry.

Crystal II of the PipX-NtcA-DNA complex was originally assigned by XDS to space group I422. From space considerations (Matthews coefficient calculations; Collaborative Computational Project Number 4, 1994), the asymmetric unit could accommodate one NtcA dimer, one DNA molecule and two PipX molecules. Initial attempts to find the phases by molecular replacement using the structures of NtcA-DNA (see above) or of PipX-NtcA (Llácer *et al*. 2010) as searching models failed with both MOLREP and PHASER. We obtained satisfactory phasing solutions by using as searching models either the NtcA dimer alone or the PipX-NtcA complex with only one PipX monomeric molecule. In all these cases, the existence of packing problems in this I422 group for reasonably accommodating a second PipX molecule suggested the possibility of merohedral twinning which would increase the apparent symmetry of the crystal, thus requiring space-group re-evaluation. To assess the twinning of the PipX-NtcA-DNA crystal complex, analyses of the intensity statistics were performed in *phenix.xtriage*, from the *PHENIX* software package (Afonine *et al*. 2018) for an I4 data set, determining the twin law governing the merohedrally twinned crystal as (*–h, k, -l*), with a twinning fraction around 0.45 (0.43 or 0.48 for H-test or the Maximum likelihood method, respectively). In fact, the use of the dataset for space group I4 instead of I422 only reduced the Rsym value from 8.5% to 7.4%, which is another indication of the nearly perfect (α=0.5) merohedral twinning of the crystal. Thus, molecular replacement in space group I4 carried out with PHASER worked with a model composed of one NtcA dimer, one DNA double helical molecule and one PipX molecule, with Matthew’s coefficient of 3.25 Å^3^ D^-1^ and 62.2% solvent content. This initial model was subjected to rigid body refinement using REFMAC, applying the twin operator (-*h*, *k*, −*l*) during each round of refinement. Then, and according to the density map, some residues were removed from the model whereas other regions were moved as rigid domains using COOT. The model was finally refined with REFMAC, using the jelly body refinement tool and applying the twin operator (-*h*, *k*, −*l*) during each round of refinement. Local non-crystallographic symmetry (NCS) restraints were automatically applied. The final refined model contained in the asymmetric unit two NtcA dimers bound to two DNA duplexes, where one NtcA dimer had two PipX molecules bound as previously reported for the PipX-NtcA dimer (Llácer *et al*. 2010). The other NtcA dimer molecule only had one PipX site canonically occupied by one PipX monomer, while the other site contacted aberrantly another PipX molecule from neighbouring NtcA2-PipX2-DNA complex.

Then we obtained a dataset for crystal I at 3.8 Å resolution. Analyses of the intensity statistics in *phenix.xtriage* revealed translational pseudosymmetry. Considering that the resulting symmetry for the initially assigned space group P21 can be defined as P1211 by using the universal Herman-Mauguin space group symbols, molecular replacement was carried out with PHASER in this space-group, using as searching assembly the PipX2-NtcA2-DNA complex of crystal II, yielding a good solution of two such assemblies in the asymmetric unit, with a Matthew’s coefficient of 2.43 Å^3^ D^-1^ and 49.5% solvent content. After rigid body refinement with REFMAC, model building in COOT was alternated with a combined refinement using real-space refinement in PHENIX (Afonine *et al*. 2018) and REFMAC using the jelly body refinement tool. In REFMAC, local non-crystallographic symmetry (NCS) restraints were automatically applied. In spite of the limited resolution, both PipX-NtcA-DNA structures (I and II) exhibit very good geometries (Table 1).

### EMSA assay

A fluorescein (Flc)-labeled 159 bp DNA fragment containing the *glnA* gene promoter was generated by PCR-amplification with Deep Vent DNA polymerase (New England Biolabs), the primer pair 5’-(Flc)CACAACCAGGAACTGAAGAC-3’ and 5’-(Flc)CGCCTGCAAGATTTCGTTAC-3’, and genomic DNA of *S. elongatus* as template. The amplified fragment, purified with GeneClean (MP Biomedicals), was used in gel retardation assays by first incubating 10 min at 32°C 30 ng of this labelled DNA, in 20 μl of a solution containing 50 mM Hepes pH 8, 3 mM MgCl2, 20% glycerol, 1 mM dithiothreitol, 25 μg/ml bovine serum albumin and 0.5 μg of poly[d(I-C)] (from Roche), and, when indicated, 100 ng of purified NtcA, either wild type or carrying the V187E mutation, and 3.2 mM 2OG. At the end of the incubation the mixtures were subjected to electrophoresis at 120V for 90 min at 4°C in native 6% polyacrylamide gels in 11 mM Tris-HCl pH 8, 11 mM boric acid and 0.25mM EDTA (low ionic strength TBE buffer). Experiments were repeated independently in duplicate. Fluorescein fluorescence was visualized using a FUJIFILM-FLA-5000 imaging system.

### Modelling of an PipX-NtcA-DNA-RNAP transcriptional activator complex

To model the interactions between RNA polymerase and NtcA-bound PipX, we superimposed our PipX-NtcA-DNA complex from crystal I onto the NtcA-TAC of *Anabaena* (PDB 8H40) (Han *et al*. 2024). Our crystal I structure comprises two PipX-NtcA-DNA complexes, with NtcA subunits labeled as A, B, D, and F. Each NtcA subunit from our complex was superimposed onto the NtcA subunit of the NtcA-TAC that is closest to the sigma subunit of RNAP. In each of the four superimpositions, the corresponding PipX coordinates were integrated to generate a PDB file. These PDB files were then analyzed using the PISA server (Krissinel *et al*. 2007) to identify interfacing residues between PipX and the RNA polymerase subunits. The results showed consistent PipX interaction sites, regardless of which NtcA subunit was superimposed. For graphical representations, the superposition of our NtcA subunit F on NtcA subunit X was used. Essentially the same results were obtained when the model was generated by superimposing on the NtcA dimer of the NtcA-TAC structure, one or the other of our two NtcA dimers in our PipX-NtcA-DNA complex of crystal I.

### Bacterial two hybrid (BATCH) assays

Strains, plasmids and oligonucleotides used for this purpose are listed in Supplementary Table S1. Cloning procedures were carried out with *E. coli* XL1-Blue, using standard techniques (Sambrook, 1989). The construction of BACTH plasmids involved, in brief, the PCR amplification of the relevant coding region, its digestion with two enzymes, and its cloning into the corresponding BACTH vectors previously cut with the same enzymes. A summary is provided in Table S2. The correctness of all the constructs was confirmed by automated dideoxy DNA sequencing.

After transformation of *E. coli* BTH101 with pairwise combination of plasmids (50 ng) and subsequent selection on ampicillin (50 µg ml^-1^) and kanamycin (40 µg ml^-1^) LB plates, five clones from each plate were inoculated into 0.5 mL of LB plus antibiotics and 0.5 mM IPTG (Isopropyl β-D-1-thiogalactopyranoside) and incubated at 30 °C for 24 hours. Interactions were assayed by dropping 3 μl of each saturated culture on M63 reporter agar plates containing 0.3 % maltose, 0.0001 % thiamine, 1 mM magnesium sulphate, 0.5 mM IPTG, and 40 µg ml^-1^ X-gal; and on MacConkey reporter plates containing 1% lactose and 0.5 mM IPTG. Reporter plates were incubated for 24 (MacConkey) or 48 hours (M63) at 30 °C and photographs were taken at 24 h intervals.

### Other methods

Protein concentrations were routinely assayed by the method of Bradford (Bradford 1976) using a commercial reagent from Bio-Rad and bovine serum albumin as a standard. Sequence alignments were carried out with Clustal Omega (Sievers & Higgins 2018), using default values. Superposition of structures was carried out with programs SSM (Krissinel *et al*. 2004) and LSQKAB [CCP4 (Collaborative Computational Project Number 4, 1994)]. Contacts between NtcA and DNA were calculated using PISA (Krissinel *et al*. 2007) or program Contacts (Collaborative Computational Project Number 4, 1994). DNA helical parameters were analysed using w3DNA (Zheng *et al*. 2009). Figures of protein structures were generated using Pymol (http://pymol.sourceforge.net/).

## Results

### The structure of the NtcA-DNA-2OG ternary complex

We determined at 3 Å resolution the crystal structure of NtcA (Table 1, Crystal 1; and Fig. 1A) bound to its target DNA, a palindrome called the NtcA box (consensus sequence, GTAN8TAC) (Luque *et al*. 1994). For the purpose of crystallization, this box was prepared as a 30 bp DNA duplex incorporating a symmetric motif derived from the first half of the corresponding box of the *S. elongatus glnA* promoter (which is strongly regulated by NtcA) (Reyes *et al*. 1997; Forcada-Nadal *et al*. 2014) and the preceding nine bases. In this way, a symmetric, synthetic, and expectedly high affinity NtcA box was prepared (see below Fig. 1D). To facilitate DNA bending (characteristic for CRP family regulators) this DNA had in each strand a break point between the GTA and TAC parts of the box, with cohesive intervening ends to maintain duplex integrity (see Material and Methods).

**Figure 1.**
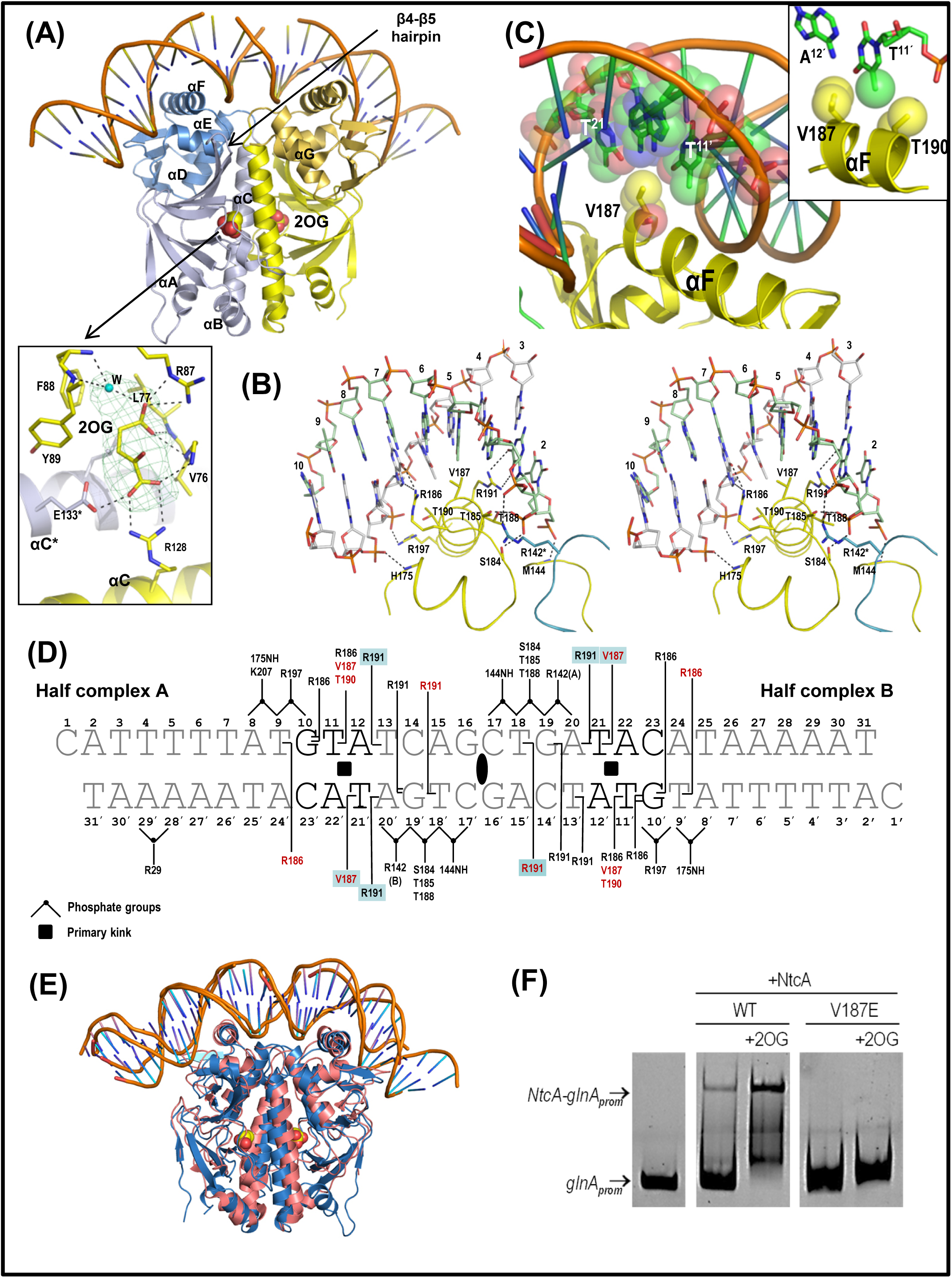
The NtcA-DNA complex structure. (*A*) Overall structure of the complex in ribbons, with bound 2OG shown in spheres representation. Some secondary structure elements of NtcA are labeled. One NtcA subunit is colored bluish and the other yellow, with the DNA binding domain shown in darker hues. The inset details the bound 2OG, modeled as in the NtcA-2OG *Anabaena* structure (Zhao *et al*. 2010), encased within its electron density omit map contoured at 2.5σ. Broken lines denote hydrogen bonds with the indicated residues. W, water. (*B*) Stereo view of the NtcA-DNA interface in one half of the NtcA-DNA complex. NtcA is colored yellow (except R142 and the loop between αC and αD from the neighbor NtcA subunit, which are bluish). One DNA chain is colored green and the other chain in gray. Residue side-chains and DNA are in sticks representation, and hydrogen bonds as black broken lines. (*C*) The DNA and helix F of NtcA to highlight the complementary Van der Waals surfaces and hydrophobic nature of the interactions of the V187 side-chain and of the bases at positions 5 and 6 of the DNA. The inset illustrates the fact that V187 and T190 clamp the methyl group of the thymine base at position 6’ of the box, thus endowing it with specificity. (*D*) Diagram of the NtcA-DNA contacts in the NtcA-DNA and PipX-NtcA-DNA (crystal I) structures. The DNA bases that characterize the consensus NtcA box are highlighted in bold-type. While the majority of the residues shown interact with DNA in both the NtcA-DNA and PipX-NtcA-DNA complexes, those shadowed in blue only interact in the complex that includes PipX. NH144 and NH175 denote that the interactions shown of amino acids 144 and 175 are mediated by the main-chain NH group of these residues. NtcA residues contributing exclusively non-polar contacts are labeled red. The interactions with the bases and with the phosphate groups are differentiated. Double interactions with a base are marked with a double line. R142(A) and R142(B) denote that the interactions are with Arg142 of the other subunit than the one providing all other contacts with the B or A moiety of the palindromic target DNA sequence. The black ellipse signals the twofold symmetry of the palindrome. The black squares mark sites of primary kinks in the DNA. *(E)* Superposition of the structures of NtcA-DNA (blue, this study) and NtcA-DNA from the NtcA-TAC complex of *Anabaena* (salmon; PDB file 8H40; Han *et al*. 2024) to show the great similarity of both structures. *(F)* Gel retardation assays of biotinylated DNA encompassing the *S. elongatus glnA* promoter without (left) or with (panels to the right) 100 ng of NtcA either wild-type or hosting the V187E mutation, in the absence or, when indicated, in the presence of 3.2 mM 2OG. For details, see Materials and Methods.

In the crystal structure (Table 1, crystal 1) one NtcA homodimer is bound to its target DNA (Fig. 1A). The DNA binding domains (DBDs) of both subunits are wrapped around by a strongly bent (∼80°) DNA duplex, with the bending largely due to sharp kinks between bases -5 and -6 and 5 and 6 on each side of the dyad axis. In this structure the protein interacts directly with 20 of the 30 base pairs of the DNA duplex (Fig. 1A-D), with all the interactions except one (with R29, a residue of the effector binding domain, EBD) being mediated by the two DNA binding domains (DBDs). The major groove of the double helix, hosting in each of two successive turns a half of the palindromic signature sequence, hosts in each turn the initial two turns of the DNA binding helix (helix F) of one subunit or the other subunit of NtcA (Fig. 1B). The axes of the helices are approximately perpendicular to the direction of the DNA chain. In this way, amino acid side chains emerging radially from each F helix can interact with the surrounding bases and phosphates in each turn of the major groove (Fig 1B, C). In addition, phosphates in the minor groove of the DNA preceding and following the consensus motif make some contacts with protein amino acids (Fig 1B, D). The NtcA dimer in this complex closely resembles the NtcA dimer reported previously in the “active” 2OG-bound conformation lacking DNA from *S. elongatus* (Llácer *et al*. 2010) or from *Anabaena* sp. (Zhao *et al*. 2010) (rmsd for the superimposition of the dimers, 0.72 Å for 421 C^α^ atoms or 0.77 Å for 388 C^α^ atoms, respectively; and see superimposition in Supplementary Fig. S1A). Indeed, our present DNA-bound structure also exhibits the effector 2OG bound on both EBDs via direct and water-mediated hydrogen bonds (inset of Fig. 1A).

Our structure of DNA-bound NtcA is also highly similar to that of the NtcA dimer contained in the recently reported cryoEM structure of NtcA-TAC complex from *Anabaena*, a complex that also includes DNA (Han *et al*. 2024) (rmsd for the superimposition of the dimers, 2.46 Å for 369 C^α^ atoms) (Fig. 1E). Actually, our NtcA-DNA complex is also quite similar to the complex of CRP with its target DNA (Supplementary Fig. S1B) (rmsd for the superimposition of 353 C^α^-atoms with the *E. coli* CRP dimer bound to DNA, 2.69 Å). The reported structures of DNA complexes of other members of the CRP-FNR family are also highly similar (Supplementary Fig. S1C) (Parkinson *et al*. 1996, Levy *et al*. 2008, Bonnet *et al*. 2013, Hall *et al*. 2016, Werel *et al*. 2023), and thus our present results add up to the existing evidence in favour that all members of the CRP transcription factor superfamily activate transcription by binding to DNA essentially in the same way.

### NtcA box recognition

The interactions between NtcA and DNA are mediated by residues located in the helix-turn-helix motif of NtcA, with the specificity being provided by residues of helix αF, which is the one that is inserted in the major groove of DNA (Fig. 1B). Nevertheless, the interactions with phosphate groups of the DNA (Fig 1B, D) should contribute to the stability of the binding of NtcA to the DNA, but, expectedly, not to the specificity of binding. Particular stabilization is likely to be obtained from ion pairs between the positively charged side chains of R142, R197 and K207 and the phosphate groups of the DNA (Fig. 1D and Table 2). Particular mention deserves such interaction between R142 and the DNA because, unlike in other members of the CRP family, in which each subunit of the transcription factor exclusively interacts with one half of the DNA consensus sequence, in the case of R142 of NtcA there is cross-interaction with the half of the NtcA box that predominantly interacts with the other subunit of NtcA. Interestingly, the position of R142 (a residue located in the dimerization helix αC) changes depending on the binding or lack of binding of 2OG to NtcA (Llácer *et al*. 2010; and see below). Only in the presence of 2OG is R142 positioned at an appropriate distance to interact with the DNA, while in the absence of 2OG the orientation of αC is such that R142 is too far to make contact with the DNA. Therefore, the indicated (Fig. 1D) interaction of R142 with the DNA phosphate appears particularly important, possibly being a key determinant for enhancing NtcA affinity for its box when 2OG is bound (see Discussion).

**Table 2.**
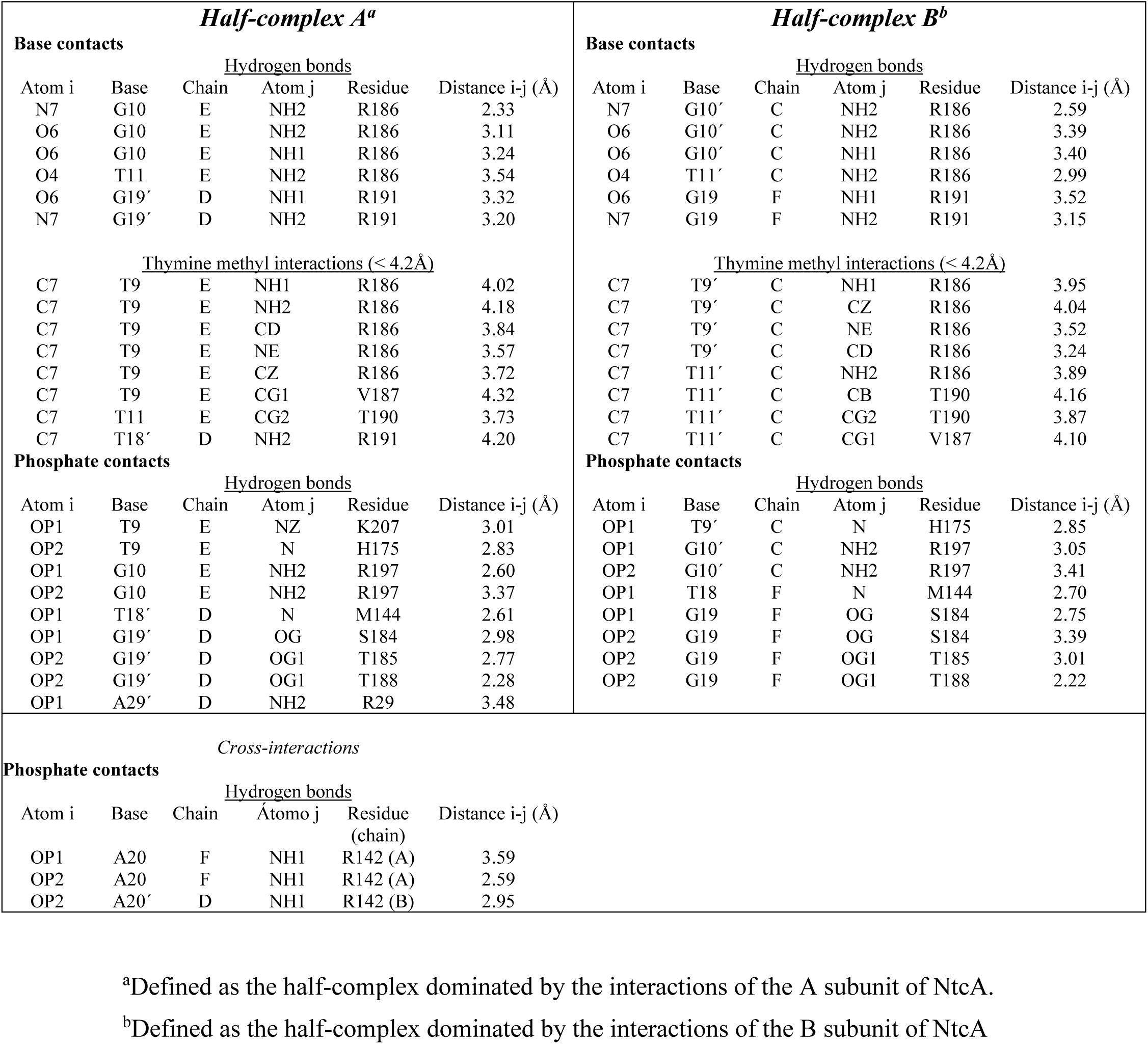
NtcA-DNA contacts.

As already mentioned, residues of helix F confer specificity to the interaction with the NtcA box, since only four residues of this helix, all of them invariant in the NtcA sequences across cyanobacterial species, R186, V187, T190, and R191, directly contact DNA bases (Table 2). In the cryoEM structure of the NtcA-TAC of *Anabaena* (Han *et al*. 2024), at least three corresponding residues, R187, V188, and R192, also contact the DNA bases, supporting the importance of these interactions. Two residues, R186 and R191 (numbering for the *S. elongatus* NtcA sequence), form hydrogen bonds with the DNA bases. Thus, the guanidinium group of R186 forms two hydrogen bonds with the O6 and N7 atoms of an invariant G:C pair in the NtcA box, at 7 base pairs from the symmetry axis of the palindrome (Figs. 1B and 1D). R191 similarly makes hydrogen bonds with the O6 and N7 atoms of a guanine at 3 base pairs from the symmetry axis, outside the conserved bases of the consensus sequence (Figs. 1B and 1D). These interactions of R191 could contribute to explain the different affinities of NtcA for different promoters that have identical consensus sequences but that show variability in the eight central bases of the palindrome (Jiang *et al*. 2000; Vázquez-Bermúdez *et al*. 2002). Massive transcriptomics data (Mitschke *et al*. 2011) identified a G at this position only in ∼50% of the putative NtcA promoters of *Anabaena sp*.

Concerning residues V187 and T190, they interact with DNA bases via van der Waals contacts with the methyl group of a thymine (T^11^ or T^11’^) at 6 base pairs from the DNA symmetry axis (Fig. 1B and 1C, and its inset). V187 is particularly notable for distinguishing NtcA from CRP, being invariant across cyanobacteria, whereas it is replaced by glutamate in CRPs. A mutation substituting V187 by glutamate (as in CRP) abolishes NtcA’s ability to bind to its target DNA, as shown in electrophoretic mobility shift assays (EMSA) (Fig. 1F). This underscores the critical role of V187 in NtcA-DNA interaction. The faint but existent retarded band in the absence of 2OG in the EMSA studies with wild-type NtcA, and the absence of this band with the V187E mutant (Fig. 1F) stresses the importance of V187 in the binding of NtcA to its DNA box even in the absence of 2OG.

### Structures of the PipX-NtcA-2OG-DNA complex

We determined the structure of the quaternary complex of NtcA with DNA, 2OG and PipX using two types of crystals (crystals I and II, corresponding to columns with numbering 2 and 3 in Table 1) grown under different conditions, and diffracting X-rays at 3.8 and 4.3 Å resolutions, respectively. Unless indicated, we will refer to the structure of crystal I (Fig. 2A-C) since it is the one obtained at the highest resolution (3.8 Å). Crystal I contained in the asymmetric unit (Fig. 2A) two complete PipX-NtcA-DNA complexes in two DNA duplexes chained by the C and G overhangs, totalizing two 30-bp-long DNA molecules hosting two NtcA dimers and four NtcA-bound PipX molecules. Despite the limited resolution, the R*factor*/R*free* values were quite good (Table 1) and, as illustrated in Fig. 2C, the map had excellent quality, allowing the tracing of most amino acid side chains, including those at the interfaces between the different components of the complex, what allowed quite detailed mapping of the interactions between the NtcA and DNA.

**Figure 2.**
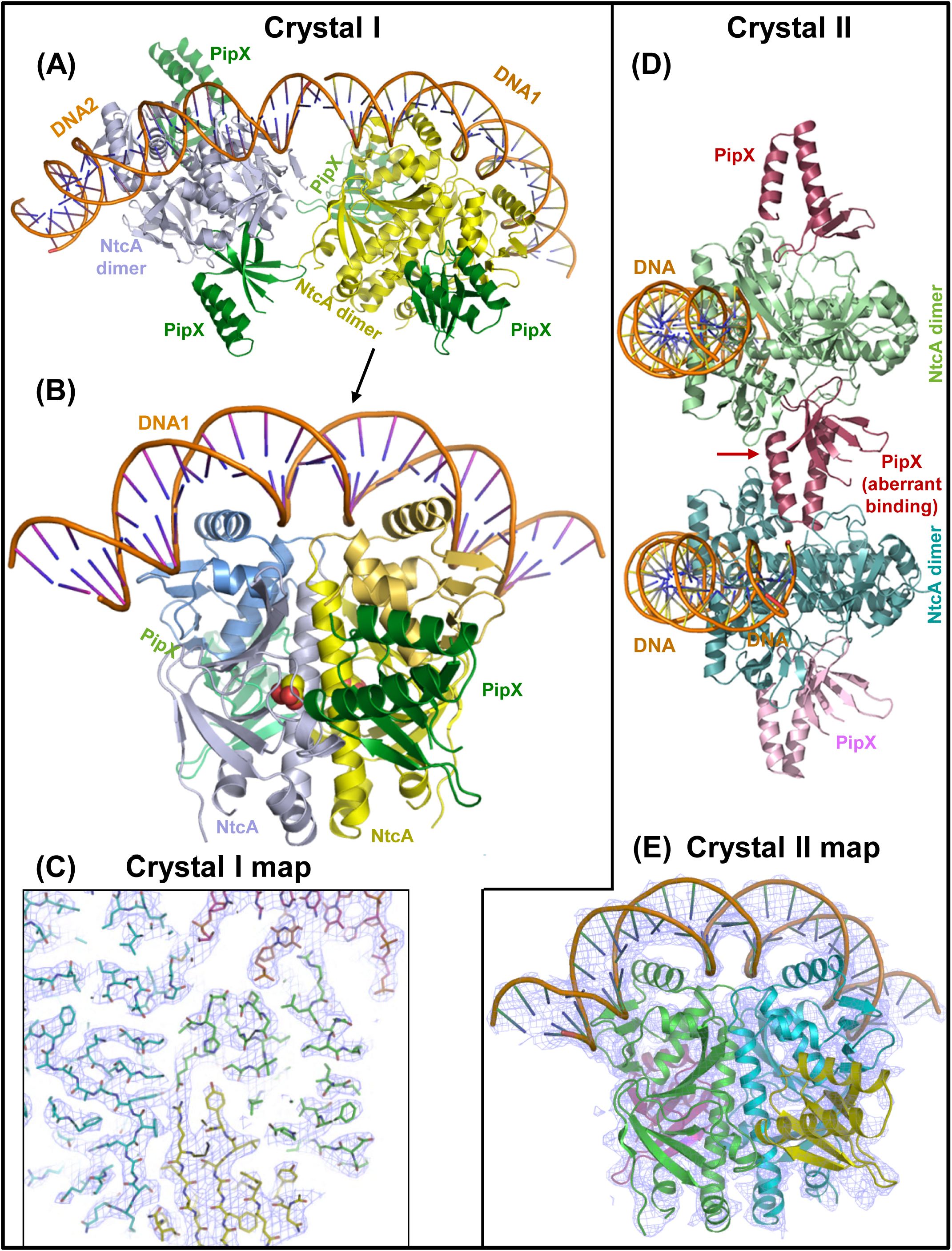
The PipX-NtcA-DNA complexes in crystals I (*A-C)* and II *(D-E),* Proteins are in ribbon representation. The different components are labelled in the same color in which they are drawn. (*A, D) C*ontents of the entire asymmetric units of the corresponding crystals. *(B)* Close-up view of the PipX-NtcA-DNA complex from the crystal at 3.8 Å resolution, with PipX in green. The allosteric effector 2OG is shown in spheres representation (C and O atoms in yellow and red, respectively). *(C)* Illustrative example of the quality of the electron density map. 2Fo-Fc map contoured at 1σ (blue-violet grid) around the sticks model for DNA (one chain in magenta, the other one in orange) and for PipX (yellow) and NtcA (one subunit cyan and the other one green). *(E)* Close-up view of one complex of crystal II to show the 2Fo-Fc electron density map at 1σ encircling the model, to illustrate the quality of the map.

The two complexes in the asymmetric unit of crystal I are very similar (rmsd of 0.97 Å for 594 superimposed C^α^ atoms). The NtcA dimer and PipX have the same topography and conformation as in the complex of these two components of *S. elongatus* in the absence of DNA (Llácer *et al*., 2010) (rmsd of 1.04±0.02 Å for 547±1 superimposed C^α^ atoms), with preservation of the PipX-NtcA interactions observed in the PipX-NtcA complex (Llácer *et al*. 2010). PipX in this complex adopts the *flexed* conformation (Fig. 2A, B) in which the C-terminal helix (helix B) runs antiparallel to the preceding helix, named A, thus forming a two-helix bundle. The flexed conformation was found also when PipX was not bound to any partner (Forcada-Nadal *et al*. 2017), whereas the B helix was extended in at least one of the three PipX molecules trapped in the complex of this protein with PII (Llácer *et al*. 2010).

DNA is bound as in the NtcA-DNA complex (Fig. 1A), and PipX does not interact with DNA (Fig. 2A). As already mentioned, the interactions of NtcA with the DNA are highly similar to those in the NtcA-2OG-DNA ternary complex, with just three extra interactions made by each NtcA subunit with each half of the palindrome (Fig. 1D, highlighted in blue; and Table 3). Because of this increase in the number of interactions, PipX might activate NtcA transcription not only by the previously proposed mechanism (Llácer *et al*. 2010) of stabilizing the “active” conformation of NtcA, but also by increasing the intrinsic affinity of the “active” NtcA dimer for its target DNA, as shown in surface plasmon resonance experiments (Forcada-Nadal *et al*. 2014).

**Table 3.** NtcA-DNA contacts in PipX-NtcA-DNA complex (crystal I)

| *Half-complex A<sup>a</sup>* |  |  |  |  |  | *Half-complex B<sup>b</sup>* |  |  |  |  |  |
| --- | --- | --- | --- | --- | --- | --- | --- | --- | --- | --- | --- |
| Base contacts |  |  |  |  |  | Base contacts |  |  |  |  |  |
| Atom i | Base | Chain | <u>Hydrogen bonds</u> |  | Distance i-j (Å) | Atom i | Base | Chain | <u>Hydrogen bonds</u> |  | Distance i-j (Å) |
|  |  |  | Atom j | Residue |  |  |  |  | Atom j | Residue |  |
| N7 | G10 | C | NH2 | R186 | 2.78 | O6 | G10' | K | NH1 | R186 | 3.39 |
| O4 | T13 | C | NH1 | R191 | 3.32 | N7 | G10' | K | NH2 | R186 | 2.57 |
| N7 | G19' | K | NH2 | R191 | 2.91 | N7 | G19 | C | NH2 | R191 | 3.10 |
| N6 | A20' | K | NH1 | R191 | 3.26 | O6 | G19 | C | NH1 | R191 | 3.50 |
| <u>Thymine methyl interactions (< 4.2Å)</u> |  |  |  |  |  | <u>Thymine methyl interactions (< 4.2Å)</u> |  |  |  |  |  |
| C7 | T9 | C | CZ | R186 | 3.71 | C7 | T9' | K | CZ | R186 | 3.69 |
| C7 | T9 | C | NH2 | R186 | 3.65 | C7 | T9' | K | NH2 | R186 | 3.73 |
| C7 | T9 | C | NE | R186 | 3.55 | C7 | T9' | K | NE | R186 | 3.48 |
| C7 | T11 | C | CG2 | T190 | 3.99 | C7 | T11' | K | CG2 | T190 | 4.02 |
| C7 | T21' | K | CG2 | V187 | 3.67 | C7 | T18 | C | CZ | R191 | 3.59 |
|  |  |  |  |  |  | C7 | T18 | C | NH2 | R191 | 3.77 |
|  |  |  |  |  |  | C7 | T18 | C | NE | R191 | 3.75 |
|  |  |  |  |  |  | C7 | T18 | C | NH1 | R191 | 3.97 |
|  |  |  |  |  |  | C7 | T21 | C | CG2 | V187 | 3.75 |
| Phosphate contacts |  |  |  |  |  | Phosphate contacts |  |  |  |  |  |
| Atom i | Base | Chain | <u>Hydrogen bonds</u> |  | Distance i-j (Å) | Atom i | Base | Chain | <u>Hydrogen bonds</u> |  | Distance i-j (Å) |
|  |  |  | Atom j | Residue |  |  |  |  | Atom j | Residue |  |
| OP1 | T18' | K | N | M144 | 3.32 | OP1 | T18 | C | N | M144 | 3.24 |
| OP1 | G19' | K | OG | S184 | 3.10 | OP1 | G19 | C | OG | S184 | 2.70 |
| OP2 | G19' | K | OG | S184 | 2.91 | OP2 | G19 | C | OG | S184 | 2.91 |
| OP2 | G19' | K | N | T185 | 3.33 | OP2 | G19 | C | N | T185 | 3.09 |
| OP2 | G19' | K | OG1 | T185 | 2.58 | OP2 | G19 | C | OG1 | T185 | 2.43 |
| OP2 | G19' | K | OG1 | T188 | 2.54 | OP2 | G19 | C | OG1 | T188 | 2.54 |
| OP1 | T9 | C | N | H175 | 3.14 | OP2 | T9' | K | N | H175 | 2.93 |
| OP1 | G10 | C | NH1 | R197 | 3.08 | OP1 | G10' | K | NH2 | R197 | 3.35 |
| OP1 | G10 | C | NH2 | R197 | 2.75 | OP2 | G10' | K | NH2 | R197 | 3.16 |
| *Cross-interactions* |  |  |  |  |  |  |  |  |  |  |  |
| Phosphate contacts |  |  |  |  |  |  |  |  |  |  |  |
| <u>Hydrogen bonds</u> |  |  |  |  |  |  |  |  |  |  |  |
| Atom i | Base | Chain | Atom j | Residue | Distance i-j (Å) |  |  |  |  |  |  |
| OP2 | A20 | C | NE | R142 (A) | 3.35 |  |  |  |  |  |  |
| OP2 | A20' | K | NH1 | R142 (B) | 3.62 |  |  |  |  |  |  |

**Table 3 (continued)**
| *Half-complex D<sup>c</sup>* |  |  |  |  |  | *Half-complex F<sup>d</sup>* |  |  |  |  |  |
| --- | --- | --- | --- | --- | --- | --- | --- | --- | --- | --- | --- |
| Base contacts |  |  |  |  |  | Base contacts |  |  |  |  |  |
| <u>Hydrogen bonds</u> |  |  |  |  |  | <u>Hydrogen bonds</u> |  |  |  |  |  |
| Atom i | Base | Chain | Atom j | Residue | Distance i-j (Å) | Atom i | Base | Chain | Atom j | Residue | Distance i-j (Å) |
| O6 | G10 | G | NH1 | R186 | 3.41 | N7 | G10' | I | NH2 | R186 | 3.33 |
| N7 | G10 | G | NH2 | R186 | 2.58 | O4 | T13' | I | NH1 | R191 | 3.41 |
| O6 | G19' | I | NH1 | R191 | 3.39 | O6 | G19 | G | NH1 | R191 | 3.27 |
| N7 | G19' | I | NH2 | R191 | 2.99 | N7 | G19 | G | NH2 | R191 | 2.78 |
|  |  |  |  |  |  | N6 | A20 | G | NH1 | R191 | 3.21 |
| <u>Thymine methyl interactions (< 4.2 Å)</u> |  |  |  |  |  | <u>Thymine methyl interactions (< 4.2 Å)</u> |  |  |  |  |  |
| C7 | T9 | G | CD | R186 | 4.20 | C7 | T9' | I | CD | R186 | 3.48 |
| C7 | T9 | G | NE | R186 | 3.48 | C7 | T9' | I | NE | R186 | 3.02 |
| C7 | T9 | G | NH2 | R186 | 3.73 | C7 | T9' | I | CD | R186 | 3.66 |
| C7 | T11 | G | CG2 | T190 | 3.44 | C7 | T9' | I | NH2 | T186 | 3.96 |
| C7 | T18' | I | NZ | R191 | 3.97 | C7 | T11' | I | CG2 | T190 | 3.90 |
| C7 | T18' | I | CZ | R191 | 3.81 | C7 | T21 | G | CG2 | V187 | 3.56 |
| C7 | T18' | I | NH1 | R191 | 4.10 |  |  |  |  |  |  |
| C7 | T18' | I | NH2 | R191 | 4.04 |  |  |  |  |  |  |
| C7 | T21' | I | CG2 | V187 | 3.75 |  |  |  |  |  |  |
| Phosphate contacts |  |  |  |  |  | Phosphate contacts |  |  |  |  |  |
| <u>Hydrogen bonds</u> |  |  |  |  |  | <u>Hydrogen bonds</u> |  |  |  |  |  |
| Atom i | Base | Chain | Atom j | Residue | Distance i-j (Å) | Atom i | Base | Chain | Atom j | Residue | Distance i-j (Å) |
| OP1 | T9 | G | N | H175 | 2.95 | OP2 | T9' | I | N | H175 | 2.87 |
| OP1 | T18' | I | N | M144 | 3.41 | OP1 | G10' | I | NH2 | R197 | 2.81 |
| OP2 | G19' | I | OG | S184 | 2.8 | OP1 | T18 | G | N | M144 | 3.33 |
| OP1 | G19' | I | OG | S184 | 3.00 | OP2 | G19 | G | OG | S184 | 2.94 |
| OP2 | G19' | I | N | T185 | 3.39 | OP1 | G19 | G | OG | S184 | 3.20 |
| OP2 | G19' | I | OG1 | T185 | 2.61 | OP2 | G19 | G | N | T185 | 3.40 |
| OP2 | G19' | I | OG1 | T188 | 2.36 | OP2 | G19 | G | OG1 | T185 | 2.63 |
|  |  |  |  |  |  | OP2 | G19 | G | OG1 | T188 | 2.51 |
| *Cross-interactions* |  |  |  |  |  |  |  |  |  |  |  |
| Phosphate contacts |  |  |  |  |  |  |  |  |  |  |  |
| <u>Hydrogen bonds</u> |  |  |  |  |  |  |  |  |  |  |  |
| Atom i | Base | Chain | Atom j | Residue | Distance i-j (Å) |  |  |  |  |  |  |
| OP2 | A20 | G | NE | R142 (D) | 3.49 |  |  |  |  |  |  |
| OP2 | A20' | I | NE | R142 (F) | 3.47 |  |  |  |  |  |  |

<sup>a</sup>Defined as the half-complex dominated by the interactions of the A subunit of the NtcA dimer composed of subunit A and B.
<sup>b</sup>Defined as the half-complex dominated by the interactions of the A subunit of the NtcA dimer composed of subunit A and B.
<sup>c</sup>Defined as the half-complex dominated by the interactions of the D subunit of the NtcA dimer composed of subunit D and F.
<sup>d</sup>Defined as the half-complex dominated by the interactions of the F subunit of the NtcA dimer composed of subunit D and F.

Although the lower resolution of the diffraction data for complex II (Fig. 2D, E) does not favour detailed structural analysis (despite the observation of quite a good map for this resolution, Fig. 2E), the PipX molecules in crystal II (Fig. 2D, E) also have their C-terminal helices flexed and do not contact the DNA, and the individual components in the complexes observed in this crystal are architecturally very similar to the corresponding elements in crystal I and in the PipX-NtcA complex having no DNA bound (Llácer *et al*. 2010). For example, rmsd values for superimposition of all the PipX molecules in the complexes in crystals I, II and in the PipX-NtcA complex without DNA (Llácer *et al*. 2010) were very low (mean of 0.62±0.09 Å for 85±1 C^α^ atoms). Furthermore, the architectures of the PipX-NtcA complexes in crystals I and II are highly similar (rmsd values of 1.00-1.09 Å for 509-594 superimposed C^α^ atoms). Nevertheless, an interesting finding in crystal II was the lack of one PipX molecule. Thus, of the two NtcA dimers bound to DNA, one has two PipX molecules bound as in the complexes observed in crystal I and in the PipX-NtcA complex (Llácer *et al*. 2010), whereas the other one has one normally bound PipX molecule while the second site for PipX is occupied by the free end of the protruding flexed helices of one of the two PipX molecules of the other dimer (Fig. 2D, labelled “aberrant binding of PipX”). Although this mode of binding may not occur in solution, the complexes in crystal II reveal that NtcA-DNA complexes with canonical (Llácer *et al*. 2010) occupation of only one PipX site might occur *in vivo*, raising the question of whether the complexes of NtcA with one PipX molecule could differ in efficiency of transcription activation relative to the complex with two PipX molecules bound. As shown below, the existence of a complex with only one molecule of PipX bound would fit the possibility that one PipX molecule could interact with RNA polymerase.

### Modelling of PipX-RNAP interactions in the NtcA-TAC complex

To explore the potential existence of interactions between PipX and RNA polymerase in the NtcA-TAC complex if PipX had been present, we superimposed our PipX-NtcA-DNA complex (Crystal I) onto the NtcA-TAC structure (PDB 8H40) (Han *et al*. 2024), following the methodology described in the Materials and Methods section. The resulting model indicated that PipX could fit into the complex without causing any steric clashes (Fig. 3A-C). The model also shows that only one PipX molecule interacts with the RNA polymerase (Fig.3A and Supplementary Fig. S2), consistent with the structural observation in our PipX-NtcA-DNA complex (crystal II), where only one PipX molecule is canonically bound to NtcA. This suggests that our observed structure of the complex with only one canonically bound PipX molecule is functionally relevant.

**Figure 3.**
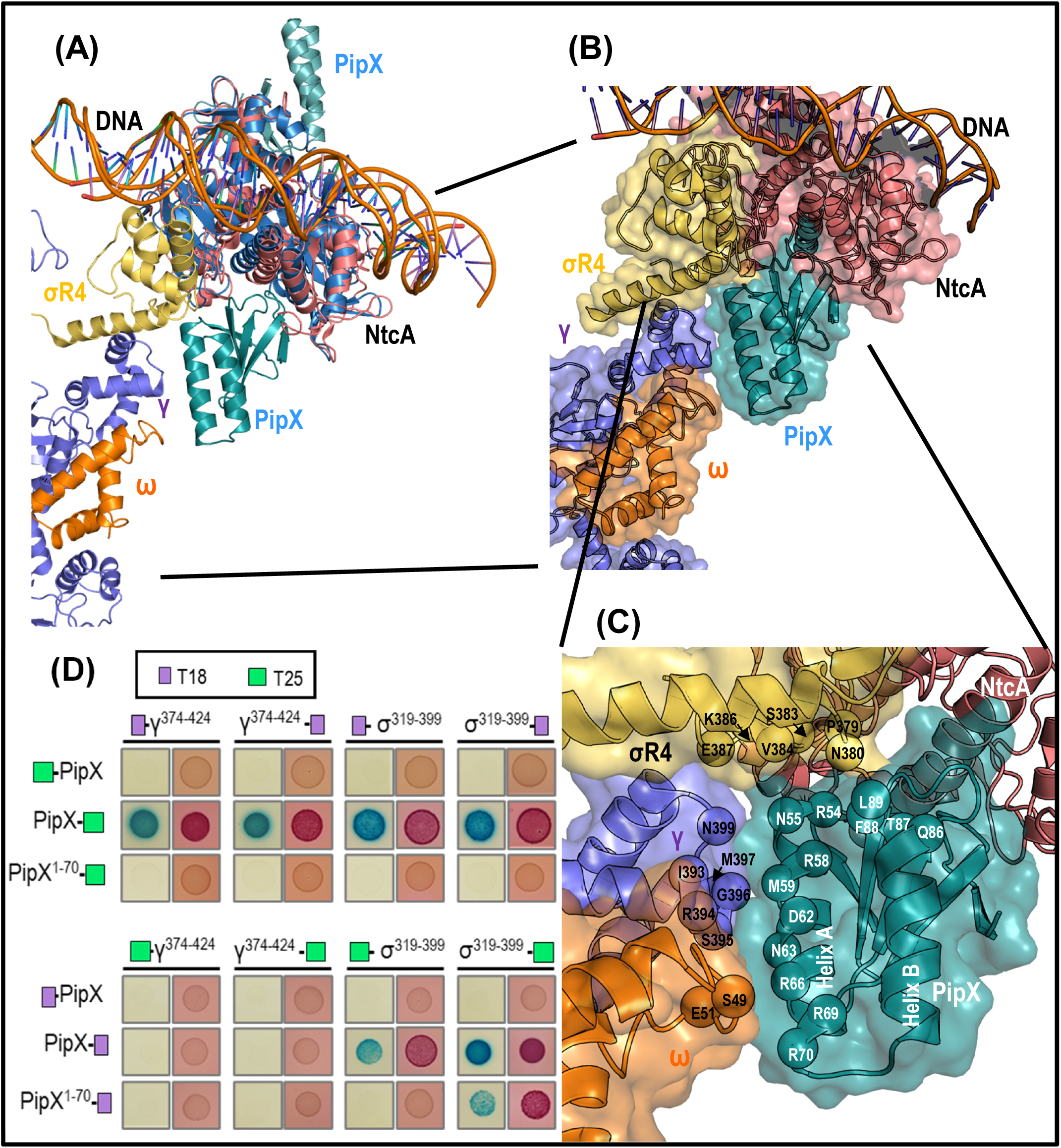
PipX-RNA Polymerase (RNAP) interactions. For clarity, only the Gamma, Omega, and domain 4 of the Sigma subunit of RNAP are depicted in all representations. *(A)* Cartoon representation of the model created by superimposing the structure of our PipX-NtcA-DNA complex from *S. elongatus* (crystal I) over the one of the NtcA-TAC complex (PDB 8H40) of *Anabaena*. The common elements of the two structures (DNA and NtcA dimer) were superimposed with Pymol (http://pymol.sourceforge.net/). DNA is in the same color for both structures, but NtcA dimers are in salmon for *Anabaena* and blue for *S. elongatus*. PipX molecules are blue-cyan and the parts shown of the σ, γ and ω subunits of RNAP are gold, blue and orange, respectively. *(B)* and *(C)* Close-ups of the NtcA-TAC complex with modelled PipX on the basis of the superimposition in (*A*). Proteins are shown in transparent surface representation to exhibit inside their chain folds in cartoon representation. In *(C)* the zoom is increased to show the PipX-interfacing residues (see Materials and Methods) and their positions within the model, alongside the corresponding RNAP interfacing residues. Amino acids are identified in single-letter code, and the two C-terminal helices A and B of PipX are labelled. (*D*) Interactions between PipX and σ^319-399^ or γ^374-424^ in the BACTH system involves α helix B. Experimental evidence using pilot bacterial two-hybrid (BACTH) analyses of PipX interactions with the indicated regions of the Gamma and Sigma subunits of RNAP (σ ^319-399^ and γ ^374-424^), which encompass the residues proposed in panel (*C*) of this figure to mediate the interactions. The relative position of the T25 or T18 domains of *CyaA-*encoded adenylate cyclase in the indicated chimeras is shown by placing the name of the indicated domain before or after the name of the potentially interacting protein. Representative photographs of plate-drop assays corresponding to the indicated pairs of fusion proteins. BTH101 co-transformants were tested in parallel on MacConkey + lactose (right) or M63 + maltose + X-gal (left) and an average of six assays performed.

The model predicts that PipX contacts through its C-terminal helices A and B with domain 4 of the sigma subunit (region 4) of RNA polymerase, as well as with the gamma subunit and, to a lesser extent, with the omega subunit (Fig. 3A-C). PISA server analysis predicts that the gamma subunit, via residues I393, R394, S395, G396, M397, and N399, could interact with helix A of PipX (potentially interacting residues of PipX, N55, R58, M59, D62, N63, and R66); that the sigma subunit region 4 could contact via its residues P379, N380, S383, V384, K386, and E387 with PipX helix A (residues R54, N55 and R58) and helix B (residues Q86, T87, F88, and L89), whereas PipX residues R66, R69, and R70 would be positioned near residues S49 and E51 of the omega subunit of the polymerase (Fig. 3C).

### Functional evidence for PipX-RNAP interactions provided by bacterial two-hybrid assays

Functional support for PipX interactions with the polymerase were obtained using bacterial two-hybrid (BACTH) assays. The BATCH assays can reveal protein-protein interactions by reconstitution in *E. coli* of the adenylate cyclase of *Bordetella pertussis* (Karimova *et al*. 1998; Battesti and Bouveret, 2012). Fragments of the sigma (σ^319-399^) and gamma (γ^374-424^) subunits predicted to interact with PipX were tested for interactions with PipX and control proteins. Since false negatives are a common issue in two-hybrid assays (Salinas *et al*. 2024), to maximize possibilities, we constructed N- and C-terminal fusions of σ^319-399^ and γ^374-424^ fragments to each of the adenylate cyclase domains (T18 and T25), generating 8 fusion derivatives that were then tested against previously validated PipX-T25, PipX-T18, T25-PipX or T18-PipX constructs (Jerez *et al*. 2021; Salinas *et al*. 2024). To support the specificity and reliability of the interactions, control proteins were also tested against σ^319-399^ and γ^374-424^ (Supplementary Fig. S3).

As shown in Fig. 3D, this technique provided clear support for the interaction *in vivo* of PipX with portions of the gamma and sigma subunits of RNAP that encompass the residues predicted by our structural modelling to participate in the interactions. In more detail, interaction signals between PipX-T25 and σ^319-399^ or γ^374-424^ derivatives were obtained in all four tested pairs, while PipX-T18 interacted with T25-σ^319-399^ or σ^319-399^-T25. None of the four polymerase fragments gave interaction signals with any of the proteins used as controls (Fig. S3), thus supporting the specificity of the interactions observed with PipX. To provide further control for the structural model, we next tested whether deleting the C-terminal helix B of PipX impaired interaction signals from positive PipX/σ^319-399^ or PipX/γ^374-424^ pairs since the prediction from the structural modelling was that three residues of helix B participate in the interactions (see above). As expected, signals were abolished or severely reduced in all cases.

Not surprisingly, false negatives were also obtained amongst some of the relevant and control proteins assayed (Fig. 3D and Fig. S3). For instance, T25-NtcA or T18-NtcA failed to interact with σ^319-399^, despite direct structural evidence (Han *et al*. 2024) or predictions (this work), suggesting that adding extra domains at the N-terminus of NtcA disrupts the interactions with the corresponding polymerase fragment. The same consideration may apply to PipX since both T25-PipX and T18-PipX gave negative results with the polymerase fragments tested. On the other hand, the failure of PipX-T18 to give interaction signals with γ^374-424^ might be attributable to stoichiometric differences, since in this system, due to differences in plasmid-copy numbers, fusions to T18 are systematically expressed at higher levels than fusions to T25 (Battesti and Bouveret, 2012). If that was the case, the implication is that comparatively more γ^374-424^ than σ^319-399^ would be required to detect interactions with PipX in our assays.

Overall, these BATCH results agree with the predictions made by the structural modelling and provides strong support for the involvement of PipX in direct contacts with specific domains of the RNA polymerase subunits sigma and gamma at NtcA regulated promoters.

### Newly identified NtcA Apo forms and NtcA Activation by 2OG

NtcA exhibits structural variability depending on its active or inactive state dictated by the presence or absence of 2-oxoglutarate (2OG). Active NtcA, either alone or complexed with PipX (Llácer *et al*. 2010) and DNA (present studies), maintains an essentially constant conformation that closely resembles that of other active Crp/Fnr superfamily members, highlighting a conserved mechanism of activation (Supplementary Fig. S4).

In contrast, the reported “inactive” forms of *Anabaena* NtcA without 2OG (Zhao *et al*. 2010) (called here form A) or of *S. elongatus* NtcA with aberrantly bound 2OG (Llácer *et al*. 2010) (called here form B) show significant mutual structural differences (RMSD of 3.67 Å for 306 Cα atoms; and see below) where A resembles more the active form than form B. By determining new crystal structures of *S. elongatus* NtcA (Table 1) in the absence of 2OG, we now report three additional inactive conformations which have no 2OG bound (called “apo forms”) (Fig. 4A). Two novel forms (A1 and A2, Fig. 4A) resemble but are not identical to the A form reported previously for *Anabaena* NtcA (Zhao *et al*. 2010) (Supplementary Fig. S5, leftmost two and one structures of the 1st and 2nd rows, respectively). These forms exclude that the A-type inactive form were specific for *Anabaena*. The third novel inactive form reported now (form B1, Fig. 4A) closely resembles the previously reported (Llácer *et al*. 2010) inactive form B (Supplementary Fig. S5, the two rightmost structures of the 1st row), but lacks the aberrantly bound 2OG found in the B form, with important differences with B in the 2OG site, particularly affecting residues 76-78, and 88, having nearly buried the empty 2OG site (Supplementary Fig. S6A). Despite their mutual similarities, forms A1 and A2 presented different crystal packing (C2221 and P212121, respectively), as was the case for form B1, which belonged to space group P6422 (Table 1).

**Figure 4.**
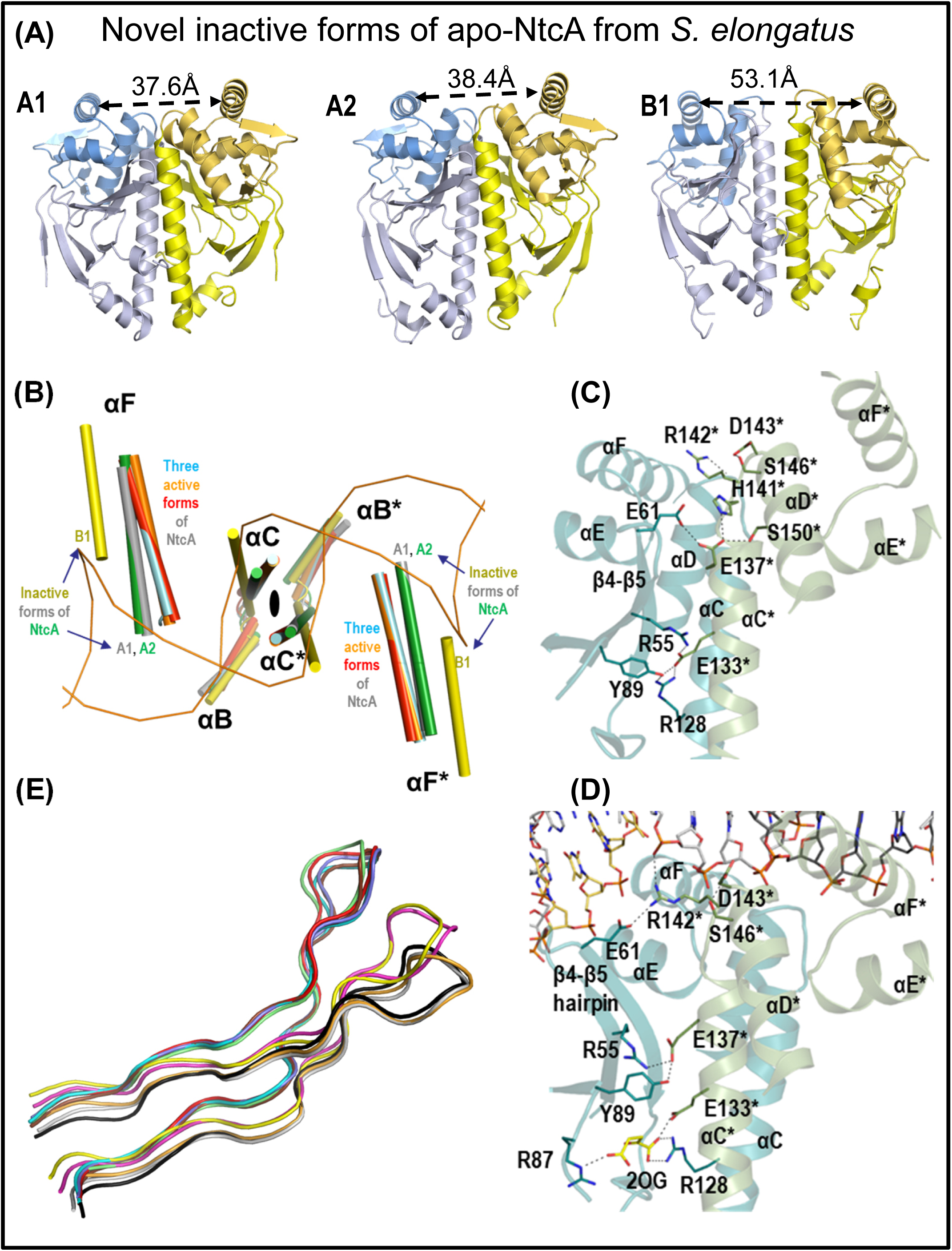
NtcA apo structures and movements of the interfacial helix, the DNA-binding helix and the β4-β5 hairpin upon activation. (*A*) Ribbon representations of NtcA in its novel apo structures, A1, A2 and B1, colored as in Fig.1A. The distance between the DNA-binding region of the F helices of both subunits are given in Å over a double-headed broken arrow. (*B*) Movements of the interfacial and the DNA-binding helices upon activation. The twofold axis of NtcA (black ellipse) is perpendicular to the paper. Only αB, the interfacial helices (αC) and the DNA-binding helices (αF) are shown as cylinders, and are labeled in the color of the indicated inactive or active form. An asterisk is used to differentiate helices from one subunit from those of the other subunit*. Active forms of NtcA:* NtcA-DNA (red), PipX-NtcA-DNA (orange), NtcA-2OG from *S. elongatus* (cyan; PDB file 2XHK). *Inactive forms of NtcA*: A1 (grey), A2 (green), B (yellow). The C and C* helices are approximately perpendicular to the paper and thus their colours are shown only at their bases. The DNA molecule belongs to the PipX-NtcA-DNA complex and is in backbone representation. *(C, D)* Mechanism of NtcA activation based on the structures of NtcA A1 *(C)* and NtcA-DNA *(D)* from *S. elongatus*. Ribbon representations showing key residues involved in signal transmission from the 2OG binding site to the DNA-binding helices. One NtcA subunit is coloured blueish and the other one greenish, with the residues of the latter subunit marked with an asterisk. Hydrogen bonds are depicted as dashed lines. An equivalent projection of inactive form A1 is shown as supplementary material (Supplementary Fig. S5B) to show that changes in essentially the same residues although of different magnitude do occur. (*C*) Comparison of β4-β5 hairpin conformation between the active and inactive forms of NtcA. NtcA-DNA (red), PipX-NtcA-DNA (brown), PipX-NtcA (cyan; PDB file 2XKO), NtcA-2OG from *S. elongatus* (blue; PDB file 2XHK), NtcA-2OG from *Anabaena* sp. (green; PDB file 3LA2), NtcA A1 (grey), NtcA A2 (black), NtcA A from *Anabaena* sp. (orange; PDB file 3LA7), NtcA B1 (yellow) and the reported NtcA B from *S. elongatus* in inactive conformation hosting aberrantly bound 2OG (magenta; PDB file 2XKP).

The comparison of the various inactive conformations of NtcA (epitomized by the structures shown in Fig. 4A) with the highly constant active conformation of this transcriptional regulator hosting correctly bound 2OG (Figs. 1A and E; 2B and E and 3A) shows that 2OG triggers a conformation exhibiting rearrangements in the EBD that propagate throughout the NtcA dimer, significantly impacting the relative positioning of the DNA-binding helices (αF) (Fig. 4B). In the active state, the distance between the DNA-interacting helices of the two subunits is short (34.8 Å), aligning perfectly for DNA binding at the two adjacent turns of the NtcA box where they sit (Figs. 1A and E and 2B and E). In contrast, these transcriptionally “inactive” forms exhibit changes of orientation and displacements of the F and F* DNA binding helices of the dimer (the asterisk signals the other subunit of the NtcA homodimer), reflected in increases in their mutual distance (to 37.6, 38.4 and 53.1 Å in A1, A2 and B1, respectively; Fig. 4A), thus preventing proper interaction of these F helices with DNA for transcriptional activation (Fig. 4B). This alteration is closely tied to shifts in the interfacial α-helices (αCs), which in active NtcA are close to the twofold axis of the dimer, forming a true coiled-coil, whereas in inactive forms they diverge more or become more parallel (Fig. 4A, B). Key to these reorientations of the C-helices are 2OG-triggered changes in the interactions across the interface that involve the side-chains of E133 and E137 (Fig. 4 C, D and Supplementary Fig. S6B).The changing partners of these two glutamate side-chains depending on whether 2OG is bound or it is not bound, favors the reorientation of the C-helices and, secondarily, of the DNA-binding domains, which become poised in a fixed conformation fitting the two turns of the major groove of the double helix in the region of the NtcA box. A critical feature in this structural modulation is the β4-β5 hairpin (Fig. 4E). In the active conformation, the interaction between 2OG and E133 disrupts prior contacts involving R55 and Y89 from the β4-β5 hairpin, freeing the latter to adopt a DNA-compatible conformation. Furthermore, this conformational change may be key to understand the second mechanism, besides increasing promoter affinity, by which 2OG could activate NtcA-dependent transcription, as seen in other family members of class II transcriptional activators such as FNR, CooA, and CRP (Bell & Busby, 1994; Leduc *et al*. 2001; Lawson *et al*. 2004). The β4-β5 hairpin belongs to the region called AR3 (“activator region 3”), the area of the transcription factor that interacts with the sigma subunit of RNA polymerase in class II TACs. Moreover, in NtcA the conformation of the β4-β5 hairpin is also important for the interaction with PipX, since in “active” NtcA, this hairpin interacts with the C-terminal end of the C-terminal helix (helix G) of the neighboring subunit (Fig. 5A), so that the conformation of this helix changes to one suitable for interacting with PipX (Llácer *et al*. 2010).

**Figure 5.**
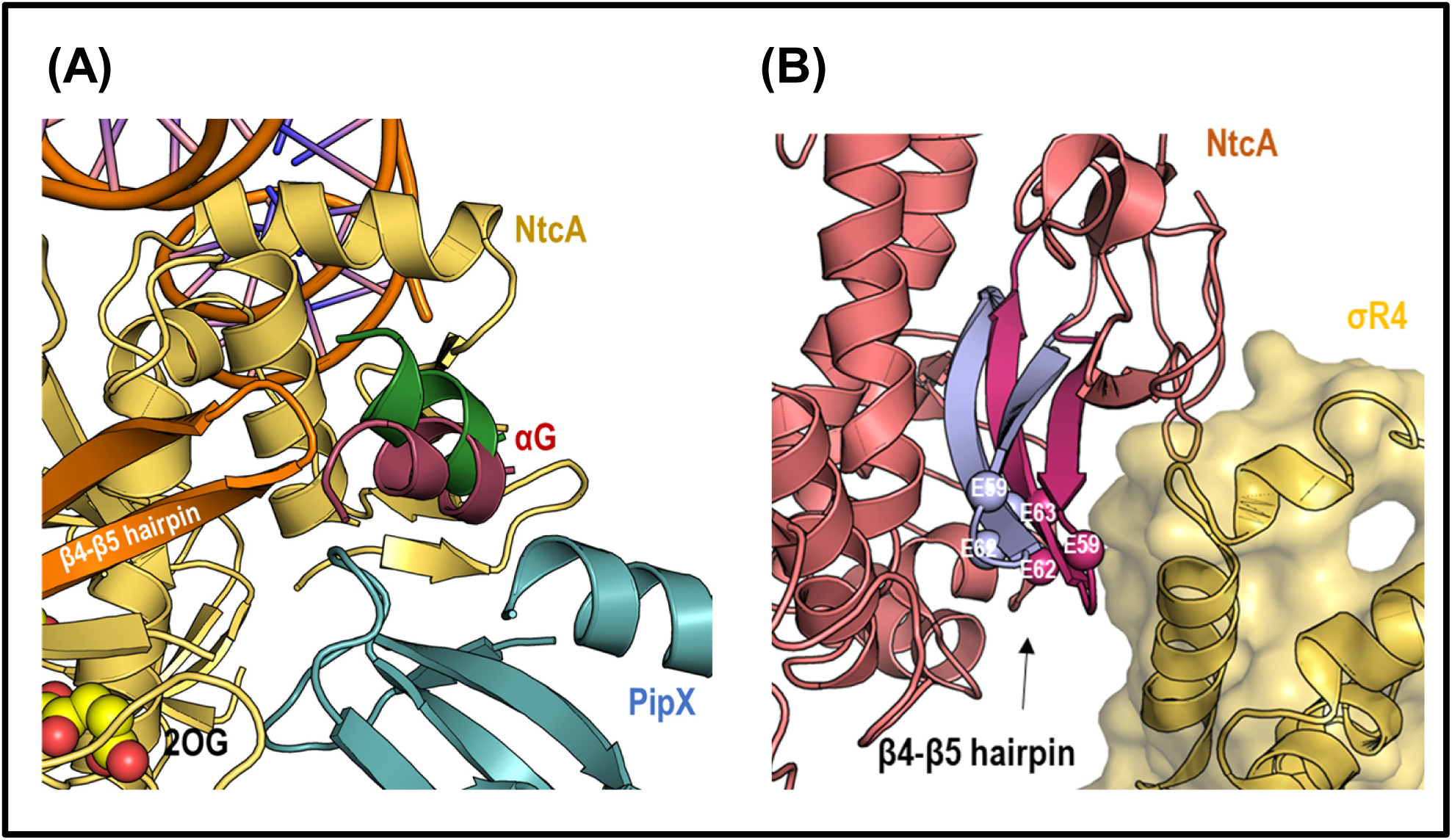
Involvement of movements of the β4-β5 hairpin and of helix G of NtcA on the activation of this transcriptional regulator by 2OG. (*A*) αG helix movement in the PipX-NtcA-DNA complex (αG helix in red) relatively to the inactive NtcA A1 structure (αG helix in green). The β4-β5 hairpin of NtcA in the PipX-NtcA-DNA complex is colored in orange. 2OG is shown in spheres representation. (*B*) Superposition of the NtcA Apo A1 structure from *S. elongatus* (this work) onto the NtcA subunit that is closest to the sigma subunit of RNAP in the NtcA-TAC complex (PDB 8H40), to highlighting the conformational change in the β4-β5 hairpin of NtcA upon 2OG binding. For clarity, only the sigma subunit domain 4 (yellow) and NtcA (salmon) of NtcA-TAC are shown, also showing (in light blue) the β4-β5 hairpin of the A1 structure, whereas the β4-β5 hairpin from the NtcA-TAC complex is hot pink. Three hairpin residues that interact with sigma R4 are shown as spheres and are labelled.

In summary, our results show that NtcA, as other factors of the CRP-FNR superfamily, can adopt multiple transcriptionally inactive conformations (Supplementary Fig. S5) in the absence of its small-molecule allosteric effector (2OG in the case of NtcA), in contrast with the constancy of the active conformation of the transcriptional regulators of this family (Supplementary Fig. S4) (Popovych *et al*. 2009; Sharma *et al*. 2009; Gallagher *et al*. 2009; Kumar *et al*. 2010; Lanzilotta *et al*. 2000; Townsend *et al*. 2014; Seok *et al*. 2014).

## Discussion

### NtcA activation and DNA binding

Transcription activation by NtcA and other members of the Crp/Fnr superfamily relies on their ability to specifically bind DNA in their active states, followed by subsequent interaction with RNA polymerase (RNAP) (Kuchinskas *et al*. 2006, Mazon *et al*. 2007, Agari *et al*. 2008, Llácer *et al*. 2010, Bonnet *et al*. 2013, Townsend *et al*. 2014, Volveda *et al*. 2015, Feng *et al*. 2016, Hall *et al*. 2016). NtcA and several related factors target class II activated promoters, where key interactions occur approximately 41.5 nucleotides upstream of the transcription start site, facilitating RNAP recruitment and the formation of active transcription complexes (Valladares *et al*. 2008, Leduc *et al*. 2001, Lawson *et al*. 2004, Bell & Busby 1994). Activation typically involves a rotation of the DBD upon induction, optimizing the arrangement of DNA recognition helices for binding (Sharma *et al*. 2009, Lanzilotta *et al*. 2000). The binding of the allosteric effector and nitrogen poorness sensor 2OG is crucial for NtcA activation, since it enables specific interactions with the NtcA box that are necessary for effective RNAP recruitment.

Recently, the determination by cryo-electron microscopy of the structure of a NtcA-TAC complex from *Anabanena* (Han *et al*. 2024) has provided insights into how DNA looping facilitates transcription activation by NtcA. DNA looping allows NtcA to bring together distantly located DNA-binding sites, enabling the transcription factor to effectively recruit RNA polymerase and activate transcription in a cooperative manner. However, the overall relatively high resolution (for cryo-electron microscopy) of the NtcA-TAC structure (3.6 Å), contrasts with the low local resolution for NtcA and the NtcA box (less than 5.5 Å), and, therefore, our present structure of NtcA bound to its target DNA at 3.0 Å resolution offers unique detail on the specific recognition of the NtcA box.

This recognition is shown here to result from a precise interaction pattern of four residues of the αF helix, Arg186, Val187, Thr190, and Arg191, with a few DNA bases of the NtcA box. These amino acid residues should allow the discrimination against other transcription factors of the same family found in bacterial cells. This discrimination was previously proven for the closely similar CRP transcription factor (Forcada-Nadal *et al*. 2014). Val187, which in CRP is replaced by a glutamate residue, is an important discriminator, since the present structures clearly predict steric and polarity clashes (not shown) in the NtcA-DNA complex if valine was replaced by glutamate. Indeed, such substitution is proven here in EMSA assays to abolish the binding of NtcA to its NtcA box. In addition, our structures also highlight the importance of extensive contacts of NtcA residues (such as Arg29, Met144, His175, Thr185, and Thr188) with phosphate groups of the target DNA. While these interactions are largely shared by CRP, the binding of Arg142 to a phosphate group is restricted to NtcA. Instead of interacting with the DNA adjacent to its subunit, Arg142 interacts with the DNA bound by the other NtcA subunit (Fig.4D). The double interaction revealed here with a DNA phosphate and with an invariant residue (Glu61) of the β4-β5 turn from the regulatory domain (EBD) of the adjacent NtcA subunit endows Arg142 with a key role in the allosteric activation of NtcA by 2OG binding.

The movement of the β4-β5 hairpin that crucially differentiates the active and inactive states of NtcA (Fig. 4E) is integral to NtcA function. In this movement, 142 is a key residue, at the center of an ion-pair network that connects the DNA with dynamic elements of NtcA like helix C and the β4-β5 hairpin (Fig. 4C, D and Supplementary Fig. S6B). The different conformations of this β hairpin depending on the activation state of NtcA (Fig. 4E) are connected to movements of the HTH DNA binding motif of the same subunit (Fig. 4D). In the inactive forms, interactions between Glu133 from one NtcA subunit and Arg55 from the other subunit, promote a different conformation of the β4-β5 hairpin that affects the HTH (Figs. 4C, D and Supplementary Fig. S6B). The β4-β5 hairpin is part of the “activator region 3” (AR3), a region interacting with the sigma subunit of RNA polymerase (Bell & Busby 1994, Leduc *et al*. 2001, Lawson *et al*. 2004). In agreement with this assignation, in the NtcA-TAC complex (Han *et al*. 2024) NtcA interacts with sigma subunit region 4 (σR4) (Fig. 3A) through residues Arg39 and glutamate residues 59, 62 and 63 (Fig. 3C), of which the later three are located in the β4-β5 hairpin (Fig. 5B). Superimposition of the inactive NtcA A1 form onto the NtcA-TAC complex highlights the shift of the position of the β4-β5 hairpin upon 2OG binding, which moves closest towards the sigma subunit of RNAP, thus strengthening the interaction of NtcA with RNAP (Fig.5B) and also illustrating why the binding of 2OG activates transcription by NtcA via promotion of effective interaction with RNAP. Interestingly, αG, the C-terminal helix characteristic of NtcA and absent in CRP mediates some of the interactions of NtcA with PipX (Llácer *et al*. 2010) and undergoes a positional shift upon 2OG binding to NtcA. This shift should facilitate NtcA interaction with PipX (Fig. 5A), likely contributing to the coactivation process.

### Roles of PipX in NtcA-dependent transcriptional activation

Determination of the PipX-NtcA-DNA complex structure has been essential in elucidating the co-activator role of PipX and it shows that NtcA and PipX, in complex with DNA retain the same conformation observed without DNA (Llácer *et al*. 2010), and it also confirms that PipX is positioned away from the DNA (Fig. 2A, B, D). Despite its moderate resolution (3.8 Å, crystal I), our present data clearly reveals additional interactions between NtcA and DNA in the PipX-NtcA-DNA complex relative to the interactions in the NtcA-DNA complex (Fig. 1D). These extra contacts, strengthening the NtcA-DNA interaction, possibly explain part (see below) of the co-activating role of PipX.

Our structure also confirms (Llácer et al., 2010) the stabilization by PipX binding of the active conformation of NtcA that is promoted by 2OG binding (Llácer *et al*. 2010). This should be another key factor in the co-activating role of PipX. By binding selectively to the active form, PipX shifts the equilibrium between inactive and active form towards the active one. Actually, each PipX molecule interacts with the EBD of one NtcA subunit and with the DNA-binding domain (DBD) of the other subunit of the dimer (Llácer *et al*. 2010), in practice stabilizing the correct orientation of this last domain and particularly of its F helix for effective DNA binding.

We previously proposed (Llácer *et al*. 2010) that PipX could also help recruit RNAP to DNA-bound NtcA, a proposal based on the fact that significant portions of the surface of PipX, particularly its two C-terminal helices, remained exposed in the PipX-NtcA complex. This exposition is now confirmed for the structure of PipX-NtcA-DNA and the interaction with RNAP further supported by our model in which we superimposed the PipX-NtcA-DNA complex structure on the NtcA-TAC lacking PipX (Han *et al*. 2024), with excellent fitting of all elements including PipX, which fits without steric clashes. Matching our model in which a single PipX molecule would be enough for helping recruit RNAP towards NtcA DNA targets, crystal II structures only had one out of the two possible PipX molecules binding to their NtcA sites (Fig. 2D). In the supercomplex, the C-terminal helices of PipX interact with region 4 of the sigma subunit of RNAP, as well as with the gamma subunit, and, to lesser extent, with the omega subunit (Fig. 3). Last but not least, BACTH interaction assays corroborated the involvement of the corresponding regions of the gamma and sigma subunits of RNAP in specific contacts with PipX.

### NtcA, a distinctive class II regulator of cyanobacteria working with a coactivator

Activation by a protein factor, in addition to an effector molecule, makes NtcA unique within the CRP/FNR family. In addition, NtcA deviates from the typical behavior of class II activators (Busby *et al*. 1999, Lawson *et al*. 2004) by not interacting with the α subunit of RNAP’s N-terminal domain (αNTD) or C-terminal domain (αCTD), as shown in the structures reported by Han et al. (2024). This raises the intriguing possibility that PipX might be necessary for these interactions to occur. It is worth noting that PipX could also fit without causing steric clashes into even larger complexes (Supplementary Fig. S7) mediating cooperative transcription activation by NtcA and NtcB (Han *et al*. 2024), a LysR-type regulator involved in nitrite utilization in *Synechococcus* (Aichi *et al*. 1997), thus suggesting a potential mechanism for combined regulation of certain genes that might require the coordinated action of two transcription factors and PipX, particularly under conditions of nitrogen depletion.

PipX is particularly important for activation of non-canonical NtcA-dependent promoters involved in nitrogen utilization like *glnN* (Espinosa *et al*. 2006, Forcada-Nadal *et al*. 2014), but it also seems that most, if not all, of the large NtcA regulon would use PipX as a coactivator (Espinosa *et al*. 2014). Therefore, coactivation by PipX appears to be globally very important for cyanobacteria, contributing to fine-tuning gene expression in response to environmental changes and to orchestrate NtcA responses to nitrogen limitation Our work provides a novel paradigm for transcriptional regulation within the CRP-FNR superfamily in bacteria and serves as a model for illustrating how co-activators like PipX can modulate transcription factor activity. These results emphasize the importance of the PipX-NtcA interaction in transcriptional regulation in cyanobacteria. The structural details elucidated here pave the way for further research into the regulatory mechanisms of other CRP-FNR family members that might involve still unrecognized novel mechanisms of coactivation. Ultimately, our findings might have applications in biotechnology and medicine, particularly in the development of strategies to control bacterial gene expression.

## Data resources

Six atomic coordinate models have been deposited in the PDB with accession codes: 9GUI, 9GUJ, 9GUK, 9GUG, 9GQU and 9GUH.

## Supplemental information

Supplemental information includes 7 figures.

## Acknowledgements.

Supported by grants of the Plan Estatal de I+D+I of the Spanish Government PID2020-116880GB-I00 to JLL; PID2020-118816GB-I00 and PID2023-149456NB-I00 to AC; and BFU2017-84264-P to VR; as well as Fundación Ramón Areces grant CIV20A6610 to VR. SB was supported by a National Grant from the Algerian Ministry of Higher Education and Scientific Research.

The authors thank T. Mata for excellent technical assistance.

## Authors’ contributions

JLL and VR conceived, supervised and guided the work in the IBV-CSIC. AF and JLL performed the structural work and drew the structural figures. AF did the EMSA studies and site-directed mutagenesis of NtcA for it. SB and PS performed BACTH experiments. AC, SB and PS prepared panel D of Fig. 3, Supplementary Tables S1 and S2 and supplementary Figure S3. JLL, VR, AC and AF wrote the paper. All the authors analyzed data, read the manuscript, made suggestions and agreed with its contents.

## Notes

**Funding** Supported by grants of the Plan Estatal de I+D+I of the Spanish Government PID2020-116880GB-I00 to JLL; PID2020-118816GB-I00 and PID2023-149456NB-I00 to AC; and BFU2017-84264-P to VR; as well as Fundación Ramón Areces grant CIV20A6610 to VR. SB was supported by a National Grant from the Algerian Ministry of Higher Education and Scientific Research.

### Competing Interest Statement

The authors have declared no competing interest.

### Summary of Updates

The revised version of the paper features a new title and includes updates on the deposition of six atomic coordinate models in the PDB, each with its respective accession codes. Additionally, the data collection and NtcA-DNA contact list tables have been updated based on the new PDB coordinates. Figure 3D has been revised with more details on the BACTH experiments, a new supplementary figure with BACTH controls has been included, and several references have been updated. The discussion section has been divided into clearer subsections to improve readability and understanding, and minor edits, especially in the discussion section, have been made to enhance the overall flow of the text. Lastly, the author contributions section has also been updated.

