## Supplementary material for "Structures of the cyanobacterial nitrogen regulators NtcA and PipX complexed to DNA fully clarify DNA binding by NtcA and recruitment of RNA polymerase by PipX"

**Supplementary Table S1.** Strains, plasmids and oligonucleotides used in BACTH assays.

| Strain | Genotype, relevant characteristics | Source or reference |
|---|---|---|
| *E. coli* XL1-Blue | *recA1 endA1 gyrA96 thi-1 hsdR17 supE44 relA1* lac [F´ *proAB* lacI$^q$ZΔ*M15* Tn*10* (Tet$^R$)]. | (Bullock *et al.* 1987) |
| *E. coli* BTH101 | F$^-$ *cya-99 araD139 galE15 galK16 rpsL1 hsdR2 mcrA1 mcrB1* | (Karimova *et al.* 2001) |
| **Plasmid** | **Description, relevant characteristics** | **Source or reference** |
| pUT18 | CyaA(225–399)T18, Ap$^R$ | (Karimova *et al.* 2001) |
| pUT18c | CyaA(225–399)T18, Ap$^R$ | (Karimova et al., 2001) |
| pKT25 | CyaA(1–224)T25, Km$^R$ | (Karimova et al., 2001) |
| pKTN25 | CyaA(1–224)T25, Km$^R$ | (Karimova *et al.* 2005) |
| pUAGC444 | T18-PipX, Ap$^R$ | (Espinosa et al., 2006) |
| pUAGC1047 | T25-PipX, Km$^R$ | (Jerez *et al.* 2021) |
| pUAGC934 | PipX-T18, Ap$^R$ | (Jerez *et al.* 2021) |
| pUAGC1045 | PipX-T25, Km$^R$ | (Jerez *et al.* 2021) |
| pUAGC1095 | PipX$^{1-70}$-T18, Ap$^R$ | (Jerez *et al.* 2021) |
| pUAGC1104 | PipX$^{1-70}$-T25, Km$^R$ | (Jerez et al., 2021) |
| pUAGC1183 | T18-$\gamma^{374-424}$, Ap$^R$ | This work |
| pUAGC1179 | T25-$\gamma^{374-424}$, Km$^R$ | This work |
| pUAGC1182 | $\gamma^{374-424}$-T18, Ap$^R$ | This work |
| pUAGC1180 | $\gamma^{374-424}$-T25, Km$^R$ | This work |
| pUAGC1193 | T18-$\sigma^{319-399}$, Ap$^R$ | This work |
| pUAGC1194 | T25-$\sigma^{319-399}$, Km$^R$ | This work |
| pUAGC1184 | $\sigma^{319-399}$-T18, Ap$^R$ | This work |
| pUAGC1181 | $\sigma^{319-399}$-T25, Km$^R$ | This work |
| **Oligonucleotide name** | **Sequence (5’— 3’)** | |
| pT25-seq | 5’ TCGGTGACCAGCGGC 3’ | |
| pUT18-sec-F | 5’ TTCACACAGGAAACAGC 3’ | |
| pUT18-sec-R | 5’ GTCGATGCGTTCGCG 3’ | |
| pKTN25-sec-R | 5’ ATGCCAGACTCCCGGTCG 3’ | |
| sigA1-BACTH-1F | 5’ GACAGGATCCCGAAGATGAAGTCGCAAAAAAC 3’ | |
| sigA1-BACTH-1R | 5’ TCAGGTACCGGGCGGATGTACTCTTTCAGGATG 3’ | |
| 1523-BACTH-1F | 5’ CTCAGGATCCCCTGCCACGGGAAATGGC 3’ | |
| 1523-BACTH-1R | 5’ GTCAGGTACCGGGCCTTCAATCACCTCTTCC 3’ | |
| 1523-BACTH-2R | 5’ GACAGGTACCCTAGCCTTCAATCACCTCTTCCAAG 3’ | |

**Supplementary Table S2.** Construction of plasmids for BACTH analysis.

| Primer Forward | Primer Reverse | Enzymes | Cloned into | Plasmid | Fusion protein expressed |
|---|---|---|---|---|---|
| sigA1-BACTH-1F | sigA1-BACTH-1R | *Bam*HI + *Kpn*I | pUT18c | pUAGC1193 | T18-$\sigma^{319\text{-}399}$ |
| | | | pKT25 | pUAGC1194 | T25-$\sigma^{319\text{-}399}$ |
| | | | pUT18 | pUAGC1184 | $\sigma^{319\text{-}399}$-T18 |
| | | | pKTN25 | pUAGC1181 | $\sigma^{319\text{-}399}$-T25 |
| 1523-BACTH-1F | 1523-BACTH-2R | *Bam*HI + *Kpn*I | pUT18c | pUAGC1183 | T18-$\gamma^{374\text{-}424}$ |
| | | | pKT25 | pUAGC1179 | T25-$\gamma^{374\text{-}424}$ |
| | 1523-BACTH-1R | *Bam*HI + *Kpn*I | pUT18 | pUAGC1182 | $\gamma^{374\text{-}424}$-T18 |
| | | | pKTN25 | pUAGC1180 | $\gamma^{374\text{-}424}$-T25 |

### Supplementary information Figure Legends

**Supplementary Figure S1.** Comparison of structures that contain “active” NtcA (A), of DNA-bound NtcA and CRP (B), and illustrations of the structures of other DNA complexes of additional members of the CRP family (C). (A) Backbone superimposition of NtcA and DNA (when present) in the structures of NtcA-2OG-DNA (yellow, present report), PipX-NtcA-2OG-DNA (cyan, present report), PipX-NtcA-2OG (magenta; PDB file 2XKO, Llácer *et al.* 2010), NtcA-2OG (green; PDB file 2XHK, Llácer *et al.* 2010), and NtcA-2OG from *Anabaena* sp. (blue; PDB file 3LA2, Zhao *et al.* 2010). (B) Stereo view of the superposition of the structures of NtcA-DNA (red, this study) and *E. coli* CRP-DNA (blue; PDB file 1J59; Parkinson *et al.* 1996). (C) Structures of other members of the Crp-Fnr superfamily, bound to DNA, to illustrate the similarities with the NtcA-DNA structure. Protein is in ribbon representation whereas ligands, when present, are in spheres representation. The structures correspond, from left to right, to PrfA (PDB 5RLS, Hall et al. 2016), $FixK_2$ (PDB 4I2O, Bonnet *et al.* 2013), CprK (PDB 3E6C; Levy *et al.* 2008) and CLR (PDB 7PZA, Werel *et al.* 2023).

**Supplementary Figure S2.** Cartoon representation of the model integrating our PipX-NtcA-DNA complex (crystal I) with the NtcA-TAC complex (PDB 8H4O) to predict potential PipX binding sites. The NtcA dimer from the *Anabaena* complex is shown in salmon, while the NtcA dimer from our PipX-NtcA-DNA complex is depicted in blue. PipX molecules are colored aquamarine. RNA polymerase subunits are represented as follows: α subunits in gray and salmon, β in pink, β' in green, γ in purple, ω in orange, and σ in yellow.

**Supplementary Figure S3.** BACTH interactions mediated by the indicated fusion proteins**.** Interactions between PipX and NtcA or PII were previously shown (Jerez *et al.* 2021; Salinas *et al.* 2024). Other details as in fig. 3D

**Supplementary Figure S4.** Structures of various members of the Crp-Fnr superfamily in active conformation. Ribbon representation. Each subunit coloured differently and labelled with its PDB file entry and the name of the protein.

**Supplementary Figure S5.** Structures of various members of the Crp-Fnr superfamily in inactive conformation. Ribbon representation. Each subunit coloured differently and labelled with its PDB file entry and the name of the protein.

**Supplementary Figure S6.** (*A*) Site for 2OG in the B1 structure and in the previously reported structure of the B inactive form of *S. elongatus* with aberrantly bound 2OG. Stereo representation. Residues involved in the interaction with 2OG are represented in sticks. B and B1 are coloured yellow and green, respectively. Black broken lines are polar contacts within the form with 2OG aberrantly bound, and red broken lines for B1 (no 2OG). Note specially the conformational change of residue F88. (*B*) Close-up of the A1 form shown similarly to the representation of Fig. 4 C for B1 forms.

**Supplementary Figure S7.** Cartoon representation of the model integrating our PipX-NtcA-DNA complex (crystal I) with the NtcA-NtcB-TAC complex (PDB 8H3V) to predict potential PipX binding sites. The NtcA dimer from the *Anabaena* complex is shown in salmon, while the NtcA dimer from our PipX-NtcA-DNA complex is depicted in blue. NtcB is colored in dark red and PipX molecules are colored aquamarine. RNA polymerase subunits are represented as follows: α subunits in gray and salmon, β in pink, β' in green, γ in purple, ω in orange, and σ in yellow

### Supplementary information References


Bonnet M, Kurz M, Mesa S, Briand C, Hennecke H, Grutter MG (2013). The structure of *Bradyrhizobium japonicum* transcription factor FixK2 unveils sites of DNA binding and oxidation. *J. Biol. Chem.* 288: 14238-14246.

Bullock, W.O., Fernandez, J.M. and Short J.M. (1987). XL1-Blue—a high-efficiency plasmid transforming *recA Escherichia coli* strain with β-galactosidase selection. *Biotechniques* 5: 376-379.

Espinosa J, Forchhammer K, Burillo S, Contreras A (2006) Interaction network in cyanobacterial nitrogen regulation: PipX, a protein that interacts in a 2-oxoglutarate dependent manner with PII and NtcA. *Mol Microbiol* 61:457-469.

Hall M, Grundström C, Begum A, Lindberg MJ, Sauer UH, Almqvist F, Johansson J, Sauer-Eriksson AE (2016). Structural basis for glutathione-mediated activation of the virulence regulatory protein PrfA in Listeria. *Proc. Natl. Acad. Sci. USA*. 113:14733-14738.

Jerez C, Salinas P, Llop A, Cantos R, Espinosa J, Labella JI, Contreras A (2021). Regulatory Connections Between the Cyanobacterial Factor PipX and the Ribosome Assembly GTPase EngA. *Front. Microbiol*. 12:781760

Karimova, G., Ullmann, A., & Ladant, D (2001). Protein-protein interaction between *Bacillus stearothermophilus* tyrosyl-tRNA synthetase subdomains revealed by a bacterial two-hybrid system. *J. Mol. Microbiol. Biotechnol. 3*: 73-82.

Karimova G, Dautin N, Ladant D (2005). Interaction network among *Escherichia coli* membrane proteins involved in cell division as revealed by bacterial two-hybrid analysis. *J. Bacteriol*. 187: 2233-43.

Levy C, Pike K, Heyes DJ, Joyce MG, Gabor K, Smidt H, van der Oost J, Leys D (2008). Molecular basis of halorespiration control by CprK, a CRP-FNR type transcriptional regulator. *Mol. Microbiol.* 70:151-167.

Llácer JL, Espinosa J, Castells MA, Contreras A, Forchhammer K, Rubio V (2010). Structural basis for the regulation of NtcA-dependent transcription by proteins PipX and PII. *Proc. Natl. Acad. Sci. USA.* 107:15397-15402.

Parkinson G, Wilson C, Gunasekera A, Ebright YW, Ebright RH, Berman HM (1996). Structure of the CAP-DNA complex at 2.5 angstroms resolution: a complete picture of the protein-DNA interface. *J. Mo.l Biol.* 260:395-408.

Werel L, Farmani N, Krol E, Serrania J, Essen LO, Becker A (2023). Structural Basis of Dual Specificity of *Sinorhizobium meliloti* Clr, a cAMP and cGMP Receptor Protein. *mBio.* 14(2):e0302822.

Zhao MX, Jiang YL, He YX, Chen YF, Teng YB, Chen Y, Zhang CC, Zhou CZ (2010). Structural basis for the allosteric control of the global transcription factor NtcA by the nitrogen starvation signal 2-oxoglutarate. *Proc. Natl. Acad. Sc.i USA* 107:12487-12492.

(A)

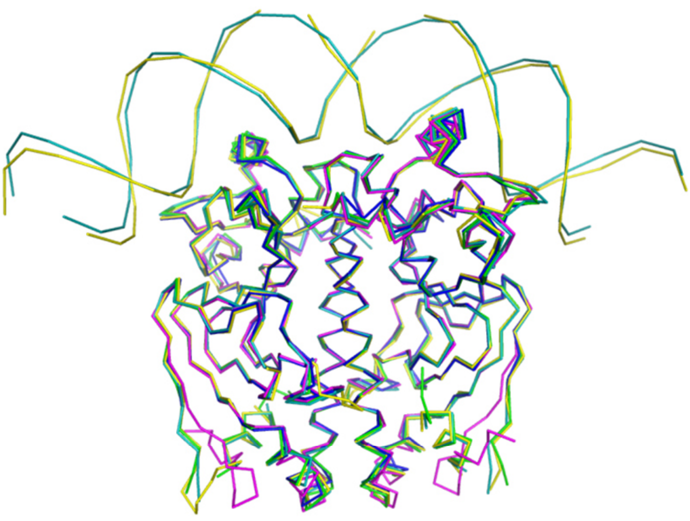

(B)

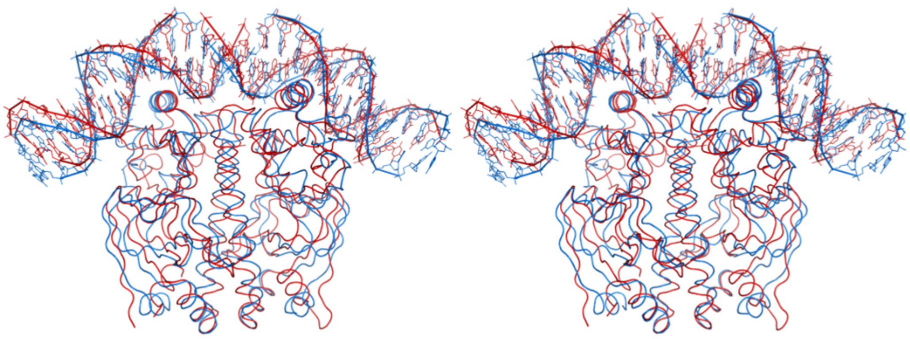

(C)

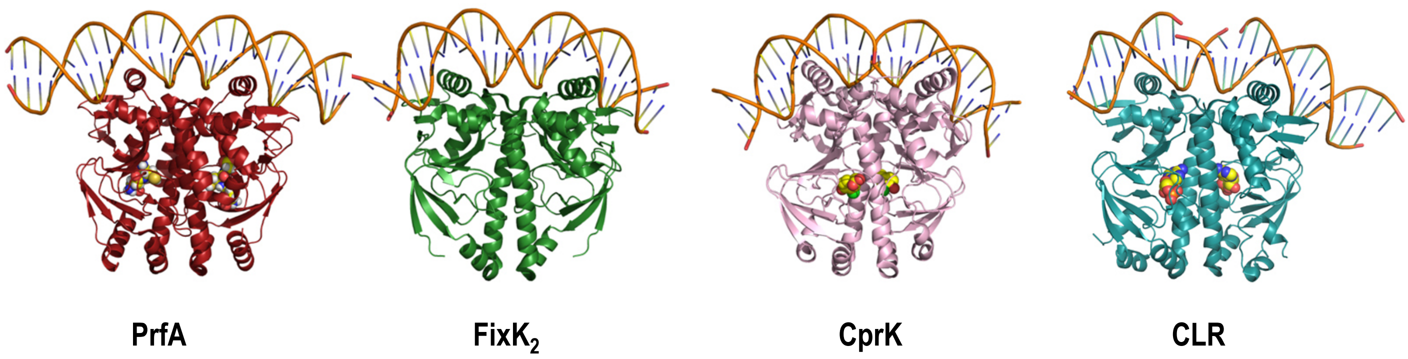


**Supplementary Fig.S1**

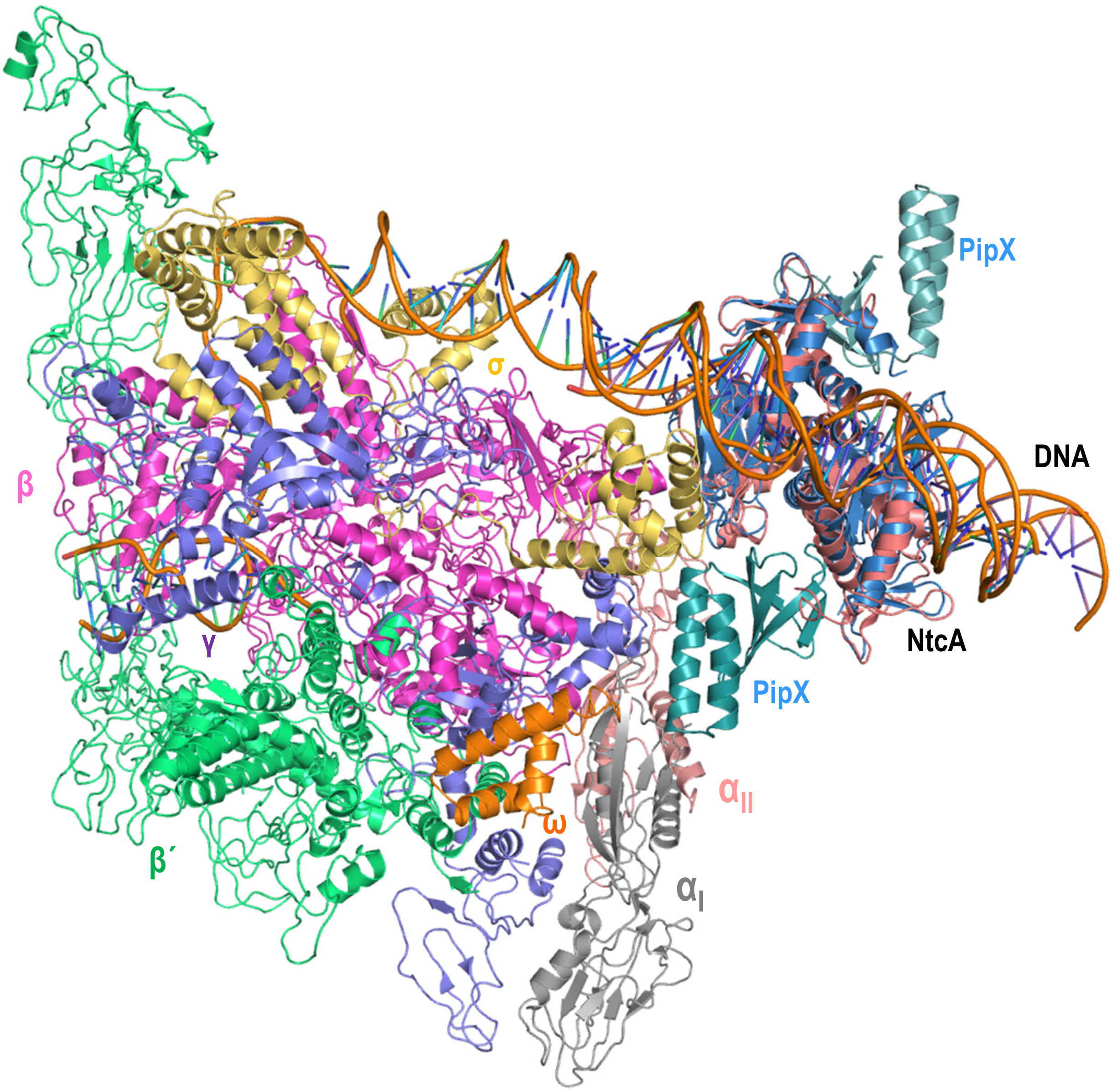


**Supplementary Fig. S2**

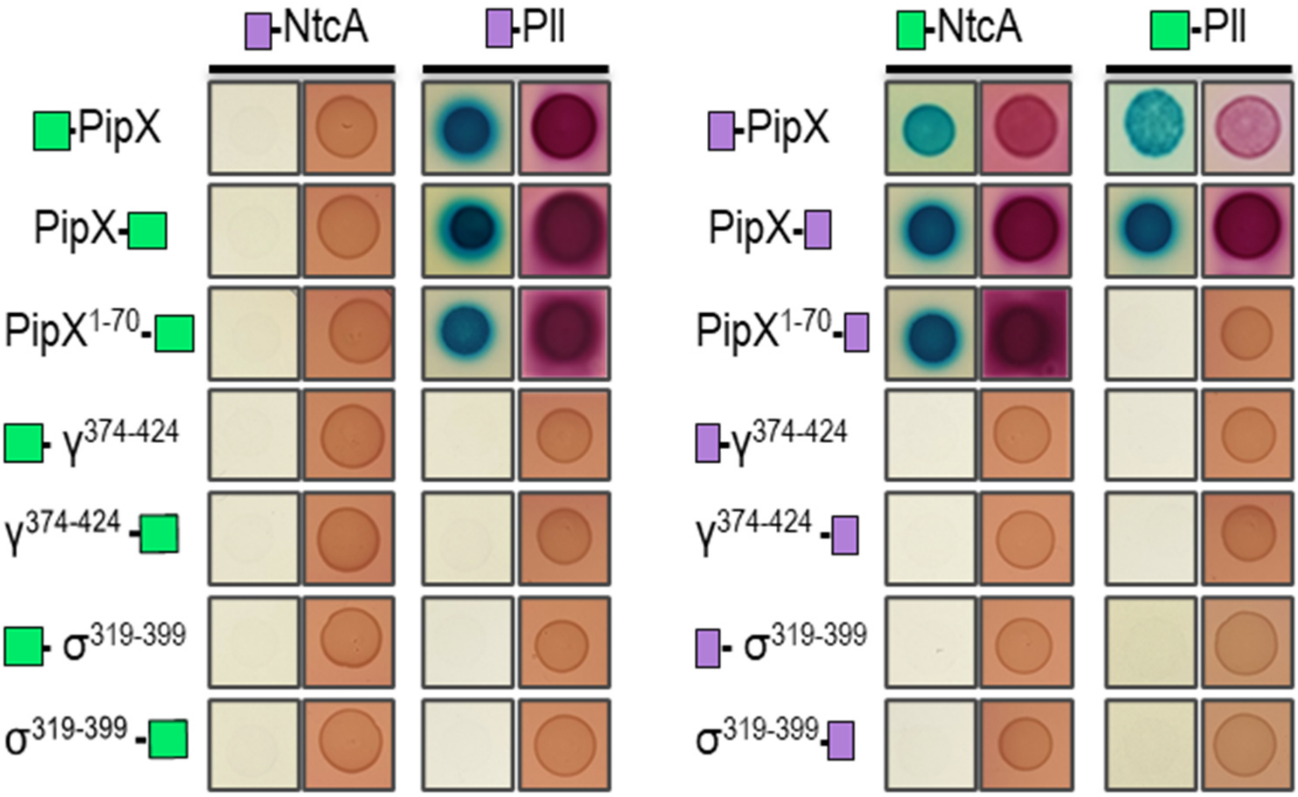


**Supplementary Fig. S3**

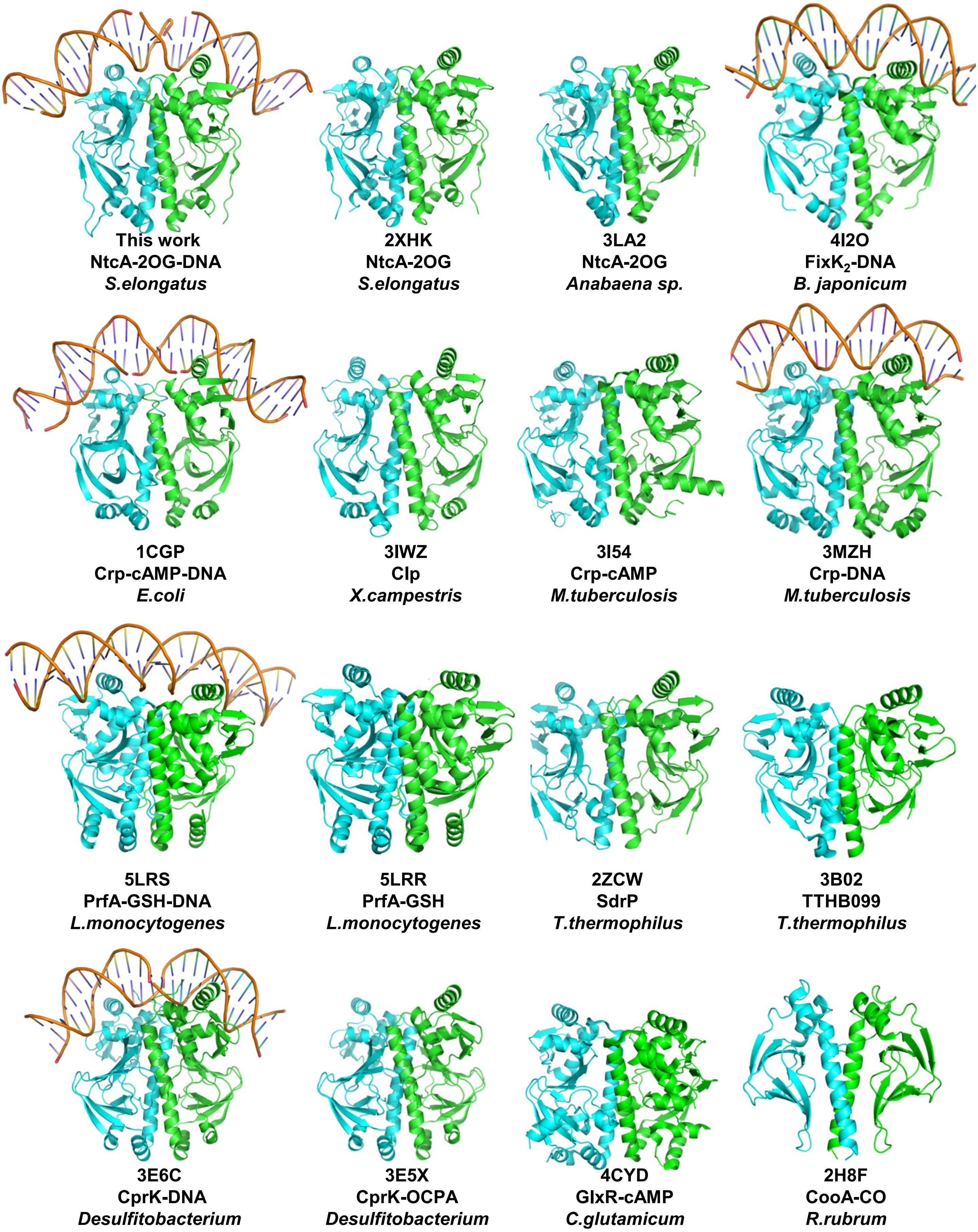


**Supplementary Fig. S4**

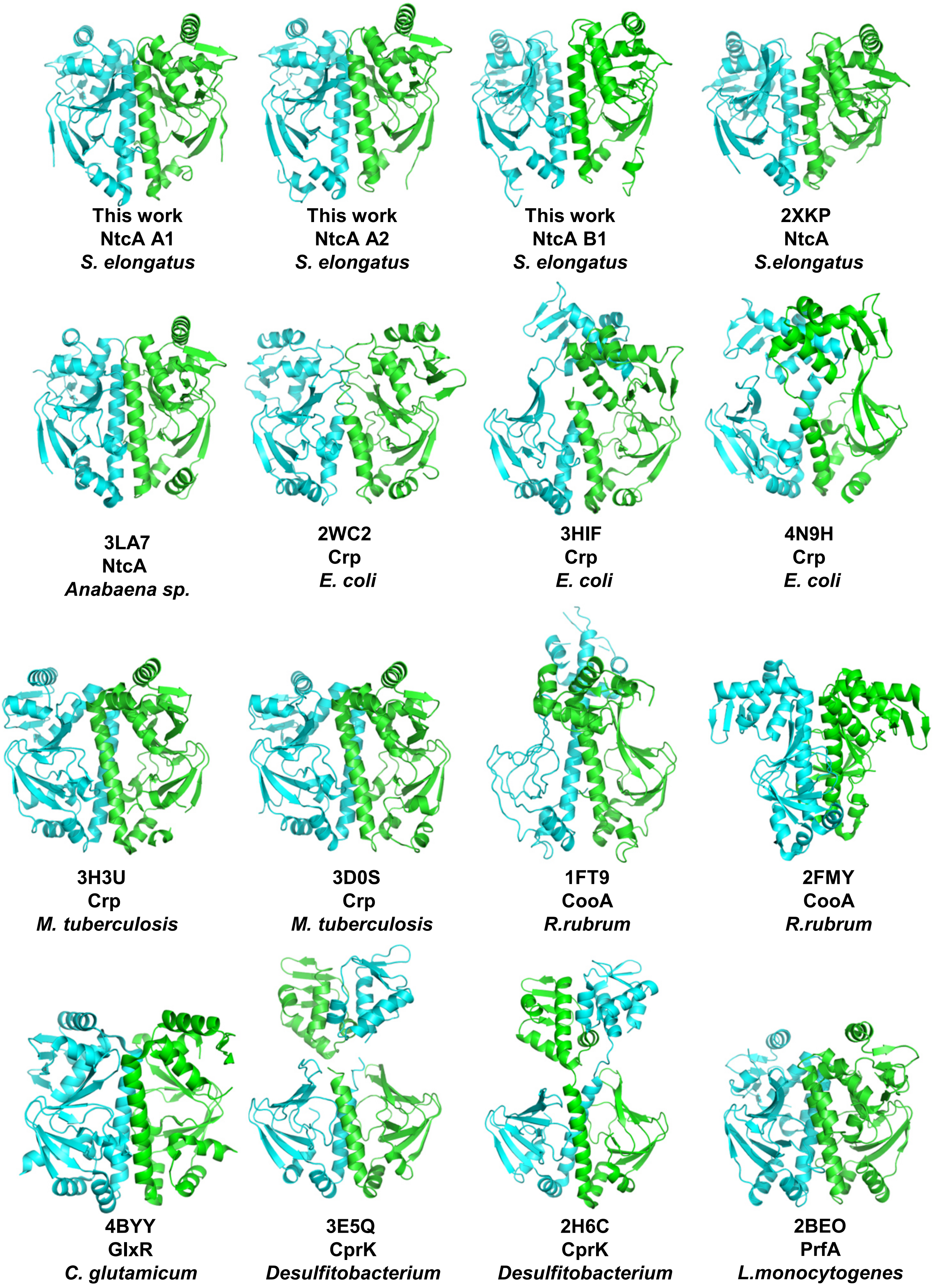


**Supplementary Fig.S5**

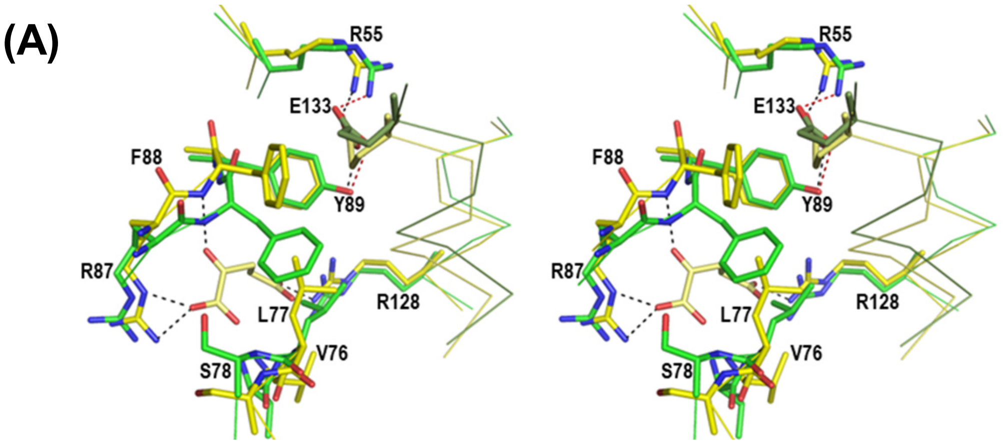


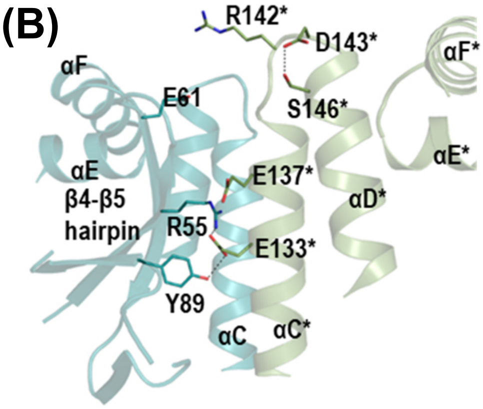


**Supplementary Fig. S6**

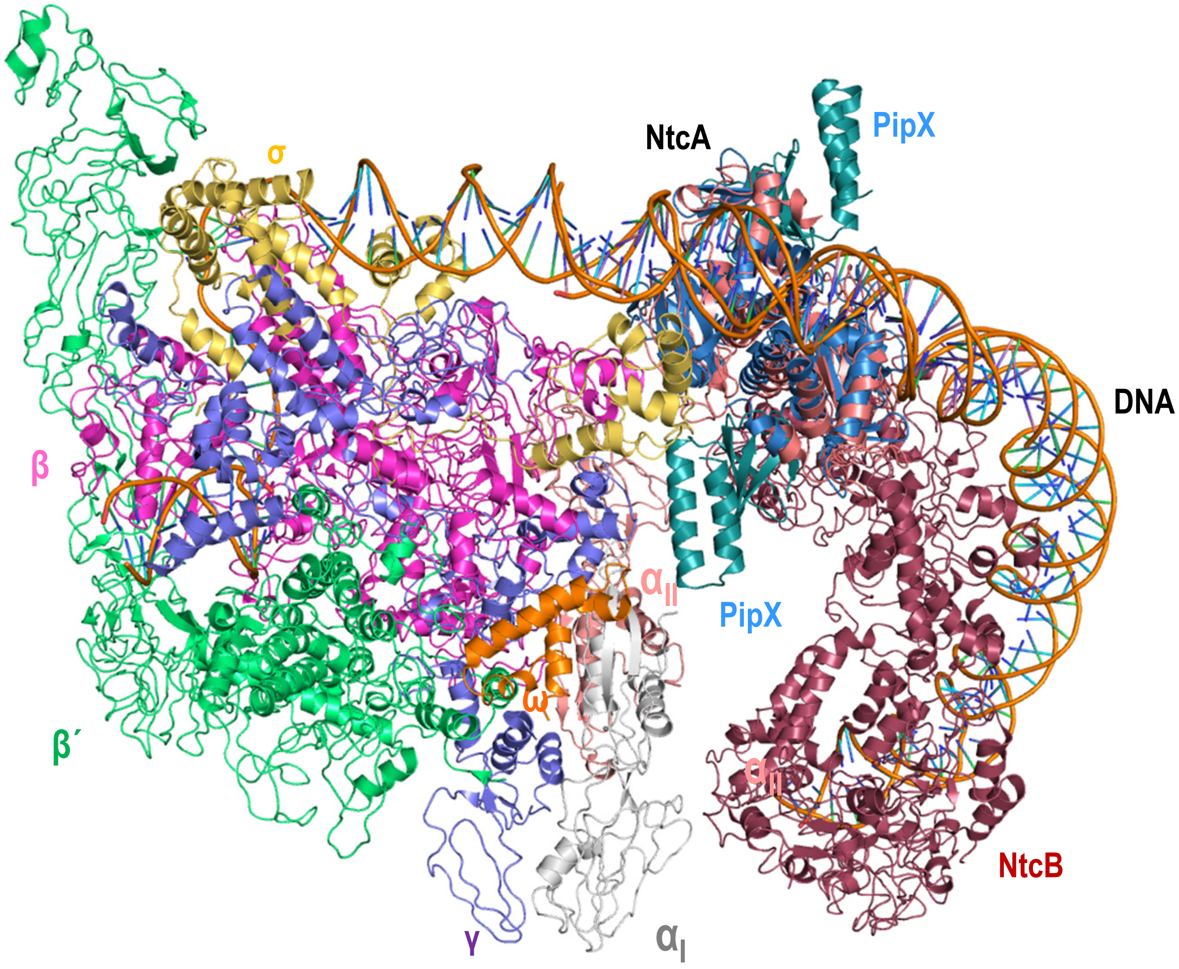


**Supplementary Fig. S7**
